# HES1 is a Critical Mediator of the SHH-GLI3 Axis in Regulating Digit Number

**DOI:** 10.1101/2020.06.17.158501

**Authors:** Deepika Sharma, Anthony J. Mirando, Abigail Leinroth, Jason T. Long, Courtney M. Karner, Matthew J. Hilton

## Abstract

Sonic Hedgehog/GLI3 signaling is critical in regulating digit number, such that Gli3-deficiency results in polydactyly and Shh-deficiency leads to digit number reductions. Anterior-posterior SHH/GLI3 signaling gradients regulate cell cycle factors controlling mesenchymal cell proliferation, while simultaneously regulating *Grem1* to coordinate BMP-induced chondrogenesis. SHH/GLI3 also coordinates the expression of additional genes, however their importance in digit formation remain unknown. Utilizing genetic and molecular approaches, we identified HES1 as a key transcriptional regulator downstream of SHH/GLI signaling capable of inducing preaxial polydactyly (PPD), required for Gli3-deficient PPD, and capable of overcoming digit number constraints of Shh-deficiency. Our data indicate that HES1, a direct SHH/GLI signaling target, induces mesenchymal cell proliferation via suppression of *Cdkn1b*, while inhibiting chondrogenic genes and the anterior autopod boundary regulator, *Pax9*. These findings fill gaps in knowledge regarding digit number and patterning, while creating a comprehensive framework for our molecular understanding of critical mediators of SHH/GLI3 signaling.

## INTRODUCTION

Development of the vertebrate limb is dependent on two signaling centers, the apical ectodermal ridge (AER) that controls proximal-distal (P-D) outgrowth and the zone of polarizing activity (ZPA), the source of *Sonic hedgehog* (*Shh*), which regulates anterior-posterior (A-P) patterning, digit number, and identity(Riddle et al. 1993; Chiang et al. 2001; Towers et al. 2008; Zhu et al. 2008). Prior to ZPA establishment within the distal posterior mesenchyme, an early A-P axis is first initiated via the expression of *Gli family zinc finger 3* (*Gli3*) and *Aristaless-like 4* (*Alx4*) within the anterior limb bud mesenchyme. GLI3 restricts the expression of *Heart and neural crest derivatives-expressed protein 2* (*Hand2*) to the posterior mesenchyme, while HAND2 antagonizes *Gli3* and *Alx4* expression(te Welscher et al. 2002a). This antagonistic relationship pre-patterns the mesenchyme allowing for the initiation of *Shh* within the posterior mesenchyme via the action of Fibroblast Growth Factors (FGFs) secreted from the AER(Charite et al. 2000; Sun et al. 2002). SHH in turn enforces *Fgf* expression (primarily *Fgf8* and *Fgf4*) within the AER, establishing a positive feedback loop that maintains both signaling centers and keeps cells within the distal mesenchyme or apical zone (AZ) in a proliferative and undifferentiated state prior to chondrogenic differentiation and digit formation(Laufer et al. 1994; Niswander et al. 1994).

During A-P patterning of the limb bud, SHH signals in a posterior to anterior gradient from the ZPA to inhibit GLI3 protein processing from the full-length activator (GLI3A) to the truncated repressor (GLI3R), thereby restricting the highest concentrations of GLI3R to the anterior limb bud mesenchyme(Wang et al. 2000). SHH-mediated regulation of GLI3 processing is critical for establishing the pentadactylous autopod with proper digit identities. Germline mutations of *Gli3* lead to multiple forms of polydactyly, and in humans is the underlying cause for Greig cephalopolysyndactyly syndrome (OMIM: 175700), Pallister-Hall syndrome (OMIM: 146510), postaxial polydactyly types A1 and B (OMIM: 174200), and preaxial polydactyly type IV (OMIM: 174700). Conventional homozygous gene deletion of *Gli3* in the mouse, represented by the *extra-toes* mutation (*Gli3*^*xt/xt*^), results in the development of generalized polydactyly (seven or greater digits without obvious identities), while heterozygous deletion or haploinsufficiency leads to the development of a single anterior extra digit or preaxial polydactyly (PPD)(Hui and Joyner 1993; Maynard et al. 2002). Conversely, germline homozygous deletion of *Shh* results in the development of only a single anterior digit with digit one identity due to the preponderance of GLI3R throughout the limb bud mesenchyme, directly implicating the SHH-GLI3 signaling axis in regulating digit number and identity(Chiang et al. 1996; Kraus et al. 2001; te Welscher et al. 2002a). Therefore a primary function of SHH within the autopod is to counteract the GLI3R-mediated constraints on digit development, and this dominance of GLI3R-mediated regulation of digit number and identity is highlighted by the indistinguishable forms of polydactyly observed in *Gli3*^*xt/xt*^ mutant and *Shh*^*−/−*^; *Gli3*^*xt/xt*^ compound mutant mice(Litingtung et al. 2002; te Welscher et al. 2002b).

Mechanistically, the SHH-GLI3 signaling axis regulates the expression of numerous factors implicated in coordinating digit number and/or identity during limb development. Unbiased gene expression studies have identified a few hundred potential candidates likely to be regulated directly or indirectly by SHH and GLI transcription factors(Vokes et al. 2008; Lewandowski et al. 2015). However, many of these genes, such as *Hoxd10-13(Knezevic et al. 1997; Zakany et al. 2004; Tarchini et al. 2006)*, *Hand2(Galli et al. 2010), Alx4(Kuijper et al. 2005), Twist1(Zhang et al. 2010; Hirsch et al. 2018), Fgf4(Lu et al. 2006), Fgf8(Moon and Capecchi 2000; Lu et al. 2006), Etv4(Zhang et al. 2009; Zhang et al. 2010), Etv5(Zhang et al. 2009; Zhang et al. 2010), Tbx2(Suzuki et al. 2004; Farin et al. 2013), Tbx3(Suzuki et al. 2004), Gata6(Kozhemyakina et al. 2014)*, and others, also in turn directly regulate *Shh/Gli3* expression and signaling that results in a feedback loop impacting digit development. Recent genetic interaction and functional studies determined that one model by which GLI3R functions to constrain digit number requires the coordinated suppression of downstream cell cycle regulators, *Cyclin d1* (*Ccnd1*) and *Cyclin dependent kinases 2, 4, and 6* (*Cdk2, Cdk4, Cdk6*), and the Bone Morphogenetic Protein (BMP) antagonist, *Gremlin (Grem1*)(Lopez-Rios et al. 2012). To control size of the limb field and digit number, this gene regulation ensures that SHH-GLI3 signaling simultaneously coordinates mesenchymal progenitor cell proliferation and the timing by which BMP signaling induces digit chondrogenesis(Lopez-Rios et al. 2012). Additional factors have been identified that potentially function downstream of the SHH-GLI3 signaling without known feedback regulation, these factors include *Paired box 9* (*Pax9*) and several NOTCH signaling pathway components, *Jagged1* (*Jag1*), *Hairy/enhancer-of-split 1* (*Hes1*), *and Hairy/enhancer-of-split related with YRPW motif protein 1* (*Hey1*)(McGlinn et al. 2005; Vokes et al. 2008; Lewandowski et al. 2015)*. Pax9* is expressed in the anterior mesenchyme of the limb bud and when deleted gives rise to a single anterior extra digit without obvious alterations to *Shh/Gli3* expression or signaling(Peters et al. 1998). In a reciprocal fashion, *Jag1, Hey1, and Hes1* are expressed in the posterior distal mesenchyme of the limb bud overlapping the ZPA (*Shh* expression) and surrounding mesenchyme(Vasiliauskas et al. 2003; McGlinn et al. 2005; Panman et al. 2006; Reinhardt et al. 2019). Genetic evidence and gene expression studies suggest that each of these genes may be regulated by SHH-GLI3 signaling, such that deletion of *Gli3* results in an anterior expansion of *Jag1, Hey1,* and *Hes1* expression with the concomitant loss of *Pax9* within the anterior mesenchyme of the limb bud(McGlinn et al. 2005; Panman et al. 2006). Further, cis-regulatory analyses identified GLI-consensus DNA binding sites and SHH-GLI3 regulation of *Pax9, Jag1,* and *Hes1(Vokes et al. 2008; Lewandowski et al. 2015)*, however it is unknown whether any of these genes are directly responsible for SHH-GLI3 function in regulating digit number or identity.

Here we define the precise role of HES1 as a critical downstream mediator of SHH/GLI3 signaling in the coordination of digit number. Using a series of genetic interaction and functional studies, we specifically demonstrate that HES1 is regulated via SHH-GLI3 signaling, is sufficient to induce PPD, is required for *Gli3*-deficient PPD, and is capable of overcoming the digit number constraints of *Shh*-deficiency. Mechanistically, we show that HES1 directly regulates mesenchymal cell proliferation, the onset of chondrogenesis, and coordinates anterior boundary formation to regulate digit number. Collectively, our data highlight a previously unknown role for HES1 in mediating SHH/GLI3 signaling during limb development.

## RESULTS

### Hes1 over-expression is sufficient to induce preaxial polydactyly (PPD)

To determine whether *Hes1* over-expression within limb bud mesenchyme is sufficient to induce PPD, we developed conditional *Hes1* gain-of-function mutant mice that carry the *Prx1Cre* transgene and *R26-Hes1*^*f/f*^ alleles(Logan et al. 2002; Kobayashi et al. 2009). *Hes1* is normally expressed within the distal posterior mesenchyme of the limb bud(Vasiliauskas et al. 2003; McGlinn et al. 2005), however utilizing this model the expression of *Hes1* is expanded throughout the limb bud mesenchyme to recapitulate the pattern of *Hes1* expression observed within *Gli3*^*xt/+*^ and *Gli3*^*xt/xt*^ limb buds(McGlinn et al. 2005).

Skeletal analyses indicate a PPD phenotype in 91% of *Prx1Cre; R26-Hes1*^*f/f*^(HES1 GOF) mutant limbs as compared to wild-type (WT) littermate controls (**Fig. 1A,B**). X-ray (**Fig. 1C**) and microCT (**Fig. 1D**) at 2-months of age reveal a sixth digit (red asterisk) anterior in position to digit 1. In approximately 9% of HES1 GOF mutants the sixth digit is an incomplete syndactylous digit, represented as 5.5 digits for quantitative purposes (**Fig. 1B**). Most other bones in HES1 GOF mice display shortening and altered morphology(Rutkowski et al. 2016), including fusions of several carpal/tarsal bones (**Fig. 1C,D and Supp. Fig. 1**). When comparing the additional anterior digit to the other digits of HES1 GOF mice, we determined that the degree of ossification at E16.5 (**Fig. 1A**) and length at 2-months of age (**Fig. 1E**) most closely resembles digit 5, suggesting that *Hes1* over-expression posteriorizes the anterior mesenchyme leading to the formation of an additional digit 5 within the anterior autopod.

**Figure 1.**
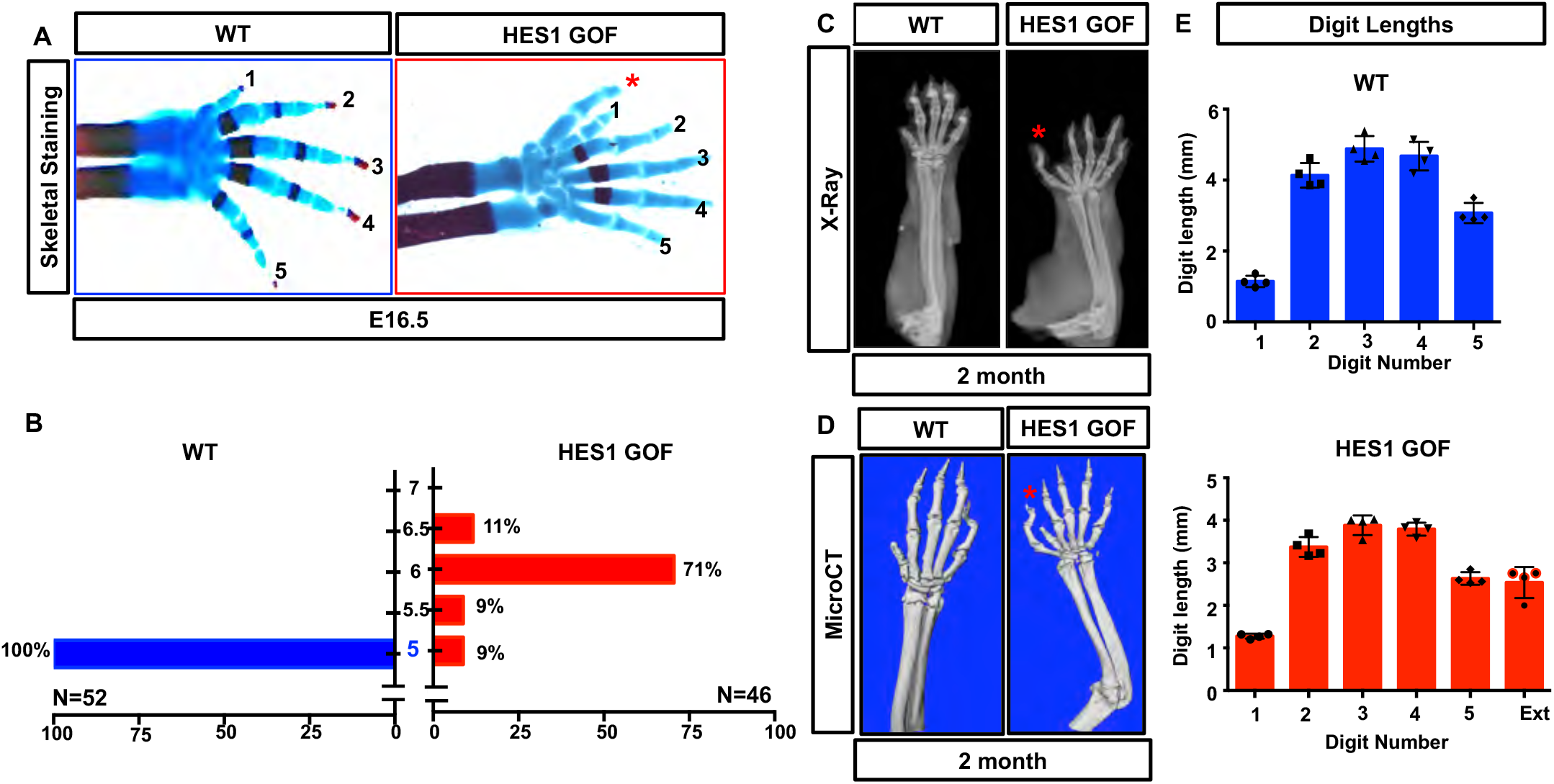
*Hes1* over-expression within limb bud mesenchyme induces preaxial polydactyly. **(A)** Alcian Blue/Alizarin Red staining of WT and *Prx1Cre;R26-Hes1*^*f/f*^(Hes1 GOF) mutant E16.5 forelimbs. Red asterisk indicates extra digit. **(B)** Quantification of digit numbers for WT (N=52) and HES1 GOF (N=46) forelimbs (χ^2^ test) (p-value <0.0001). Intervals of 0.5 represent syndactylous digits. **(C-D)** Representative X-ray and MicroCT images of 2-month old WT and HES1 GOF forelimbs. Red asterisks indicate the extra digit. **(E)** Digit length analysis of WT and HES1 GOF at 2-months of age (N=4).

To determine the regional effects of *Hes1* over-expression on digit development, we generated *ShhCre; R26-Hes1*^*f/f*^ mutant and controls. *ShhCre* activation induces *Hes1/Gfp* over-expression within posterior mesenchymal cells of the ZPA and their descendants that give rise to digits 4 and 5 (**Supp. Fig. 2A**)(Harfe et al. 2004). Skeletal staining demonstrates that over-expression of *Hes1* within the posterior mesenchyme does not induce additional digit formation (**Supp. Fig. 2B**), further suggesting that HES1 action within the anterior mesenchyme drives the PPD phenotype in HES1 GOF mice.

### *Hes1* over-expression within limb bud mesenchyme alters cell cycle kinetics

To determine potential mechanisms underlying HES1-induced PPD, we analyzed mesenchymal cell proliferation utilizing BrdU and phospho-histone H3 (PH3) immunohistochemistry/immunofluorescent (IHC/IF) staining. HES1 GOF limb buds display a significant increase in BrdU and PH3 positive mesenchymal cells within the apical zone, along the P-D axis, and within the anterior mesenchyme (**Fig. 2A,B** and **Supp. Fig. 3A,B**). Western analyses for cell cycle regulators indicate a decrease in the cell cycle inhibitor, p27, and an increase in CYCLIN D1 (CCND1) (**Fig. 2C**) within the mesenchyme of HES1 GOF limb buds. Real-time quantitative PCR (qPCR) analyses using RNA isolated from the anterior halves of E11.5 limb buds demonstrate a mild increase in *Cdk2, Cdk4,* and *Cdk6* expression and confirmed the significant decrease in *Cdkn1b* expression, encoding p27 (**Fig. 2D**). Since HES1 primarily acts as a transcriptional repressor(Ohtsuka et al. 1999), we searched for and identified putative HES1 binding sites (one E-box and two N-boxes) within 3kb of the *Cdkn1b* transcriptional start site (**Supp. Fig. 4A**). Chromatin immunoprecipitation (ChIP) using HES1 antibodies on DNA isolated from E11.5 HES1 GOF and WT limb buds demonstrated that HES1 is capable of binding the conserved E-box located approximately 200bp upstream of the *Cdkn1b* transcriptional start site (**Fig. 2E**). These data indicate that over-expression of *Hes1* enhances mesenchymal cell proliferation during limb development in part via the direct suppression of *Cdkn1b*.

**Figure 2.**
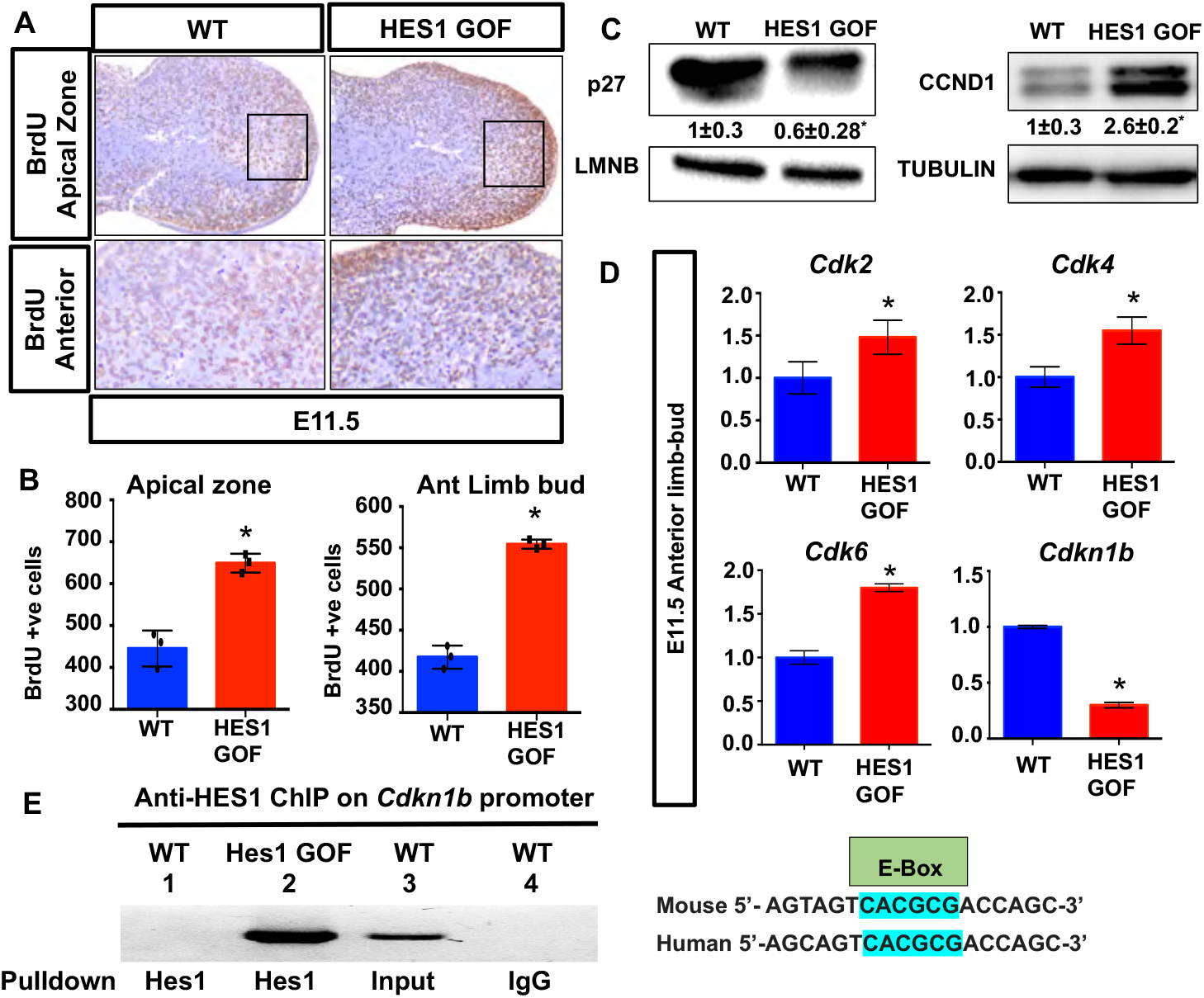
*Hes1* over-expression within limb bud mesenchyme promotes cell proliferation. **(A)** BrdU labeling of WT and HES1 GOF limb buds at E11.5 (N=3) **(B)** Quantification of BrdU positive cells within the apical zone and anterior domain (N=3). Asterisks indicate significance with a p-value < 0.05 (Student’s t-test). **(C)** Western blots of p27 and CCND1 utilizing protein isolated from E11.5 WT and HES1 GOF limb buds. LMNB and TUBULIN are loading controls. Quantification represented as Fold change± SD relative to WT. **(D)** qPCR for *Cdk2, Cdk4, Cdk6* and *Cdkn1b* using RNA isolated from the anterior halves of WT and HES1 GOF limb buds at E11.5 (N=4). Asterisks indicate significance with a p-value < 0.05 (Student’s t-test). **(E)** Schematic of conserved E-Box (HES1 binding site) within human and mouse *Cdkn1b* promoters. ChIP and PCR amplification of chromatin containing E-Box within *Cdkn1b* promoter (N=3). Lane 1 = amplification of WT chromatin pulled down with anti-HES1; Lane 2 = amplification of Hes1 GOF chromatin pulled down with anti-HES1; Lane 3 = amplification of input control chromatin (positive control); Lane 4 = amplification of WT chromatin pulled down with anti-IgG (negative control).

### *Hes1* over-expression within limb bud mesenchyme delays chondrogenesis

To further investigate the effects of *Hes1* over-expression on limb development, we generated HES1 GOF and controls from E11.5 to E12.5 and performed whole-mount *in situ* hybridization (WISH), histology, IHC, and qPCR. WISH for *Sry-box 9* (*Sox9*), a transcription factor critical for forming mesenchymal condensations and chondrocyte differentiation(Ng et al. 1997), demonstrates a modest reduction in *Sox9* expression within E11.5 HES1 GOF limb buds (**Fig. 3A**). Alcian blue staining of E12.5 HES1 GOF and WT limb bud sections demonstrates a delay in digit chondrogenesis of HES1 GOF embryos (**Fig. 3A**). IHC analyses for COL2A1 confirm the delay in cartilage formation of HES1 GOF autopods, however proximal limb condensations eventually undergo chondrogenic differentiation leading to the formation of smaller cartilage elements (**Fig. 3A**). qPCR performed on RNA extracted from the anterior halves or whole limb buds at E11.5 and E12.5 demonstrate a reduction in *Sox9, Sry-box 5* (*Sox5*), *Sry-box 6* (*Sox6*), *Type II Collagen* (*Col2a1*), and *Aggrecan* (*Acan*) expression in HES1 GOF limb buds at each time point (**Fig. 3B and Supp. Fig. 5**). Consistent with published reports(Grogan et al. 2008), our sequence analyses and ChIP studies demonstrate that HES1 is capable of binding a conserved N-box within the *Col2a1* enhancer approximately 1700bp downstream of exon 1 (**Fig. 3C and Supp. Fig 4B**). These data indicate that over-expression of *Hes1* within limb mesenchyme delays the formation of cartilage condensations and chondrogenesis within the limb skeleton, likely via direct negative regulation of chondrogenic genes such as *Col2a1* and potentially others(Grogan et al. 2008; Rutkowski et al. 2016).

**Figure 3.**
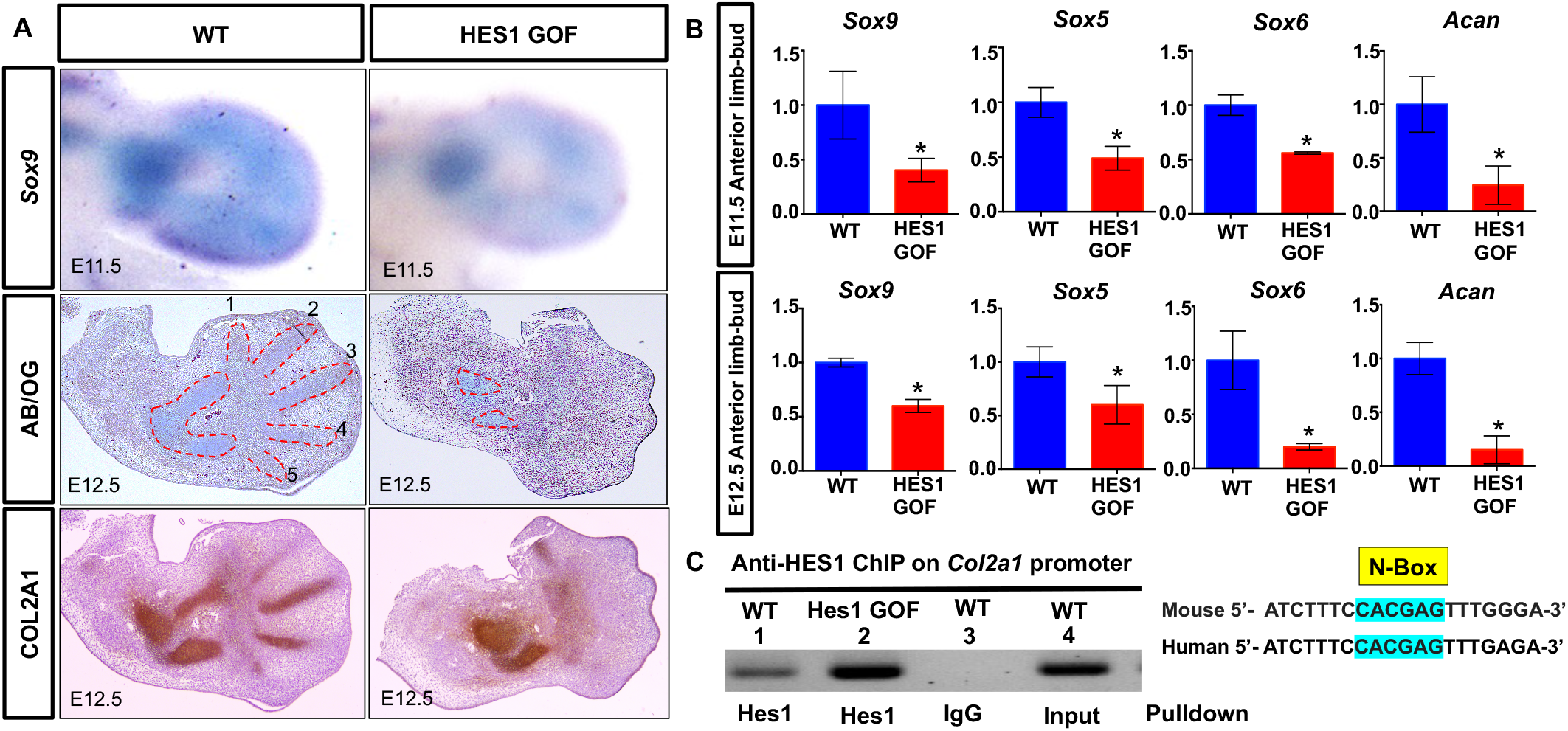
*Hes1* over-expression within limb bud mesenchyme delays chondrogenesis. **(A)** WISH for Sox9 on WT and HES1 GOF E11.5 limb buds. ABH/OG staining of WT and HES1 GOF E12.5 limb bud sections. COL2A1 IHC staining of WT and HES1 GOF E12.5 limb bud sections. **(B)** qPCR for *Sox9, Sox5, Sox9, and Acan* utilizing RNA from anterior limb bud halves of E11.5 and E12.5 WT and HES1 GOF embryos (N= 4; p-value < 0.05; Student’s t-test) **(C)** Schematic of N-Box within the human and mouse enhancer sequence of *Col2a1* located between exons 1 and 2. ChIP and PCR amplification of chromatin containing N-Box within *Col2a1* enhancer (N=3). Lane 1 = amplification of WT chromatin pulled down with anti-HES1; Lane 2 = amplification of Hes1 GOF chromatin pulled down with anti-HES1; Lane 3 = amplification of WT chromatin pulled down with anti-IgG (negative control); Lane 4 = amplification of input control chromatin (positive control).

### *Hes1* over-expression alters several SHH/GLI-associated factors critical for digit number and patterning

To assess HES1 regulation of genes implicated in PPD, we performed WISH and qPCR for known regulators of digit number and patterning, including *Pax9*, *Alx4, Fgf8* and *Bmp4*. We observed patterns of expression similar to *Gli3*^*xt/+*^ and *Gli3*^*xt/xt*^ mutant limb buds(McGlinn et al. 2005; Hill et al. 2009; Lopez-Rios et al. 2012), including decreased *Pax9* expression (white arrow), restricted *Alx4* expression to the proximal mesenchyme (red arrow), expanded *Fgf8* expression within the AER (yellow arrow), and reduced mesenchymal expression of *Bmp4* (black arrows) (**Fig. 4A**) and confirmed these results via qPCR (**Fig 4B**). These data indicate that HES1 overexpression alters the expression of several critical regulators of digit number and patterning, similar to that observed in *Gli3*-deficient polydactylies.

**Figure 4.**
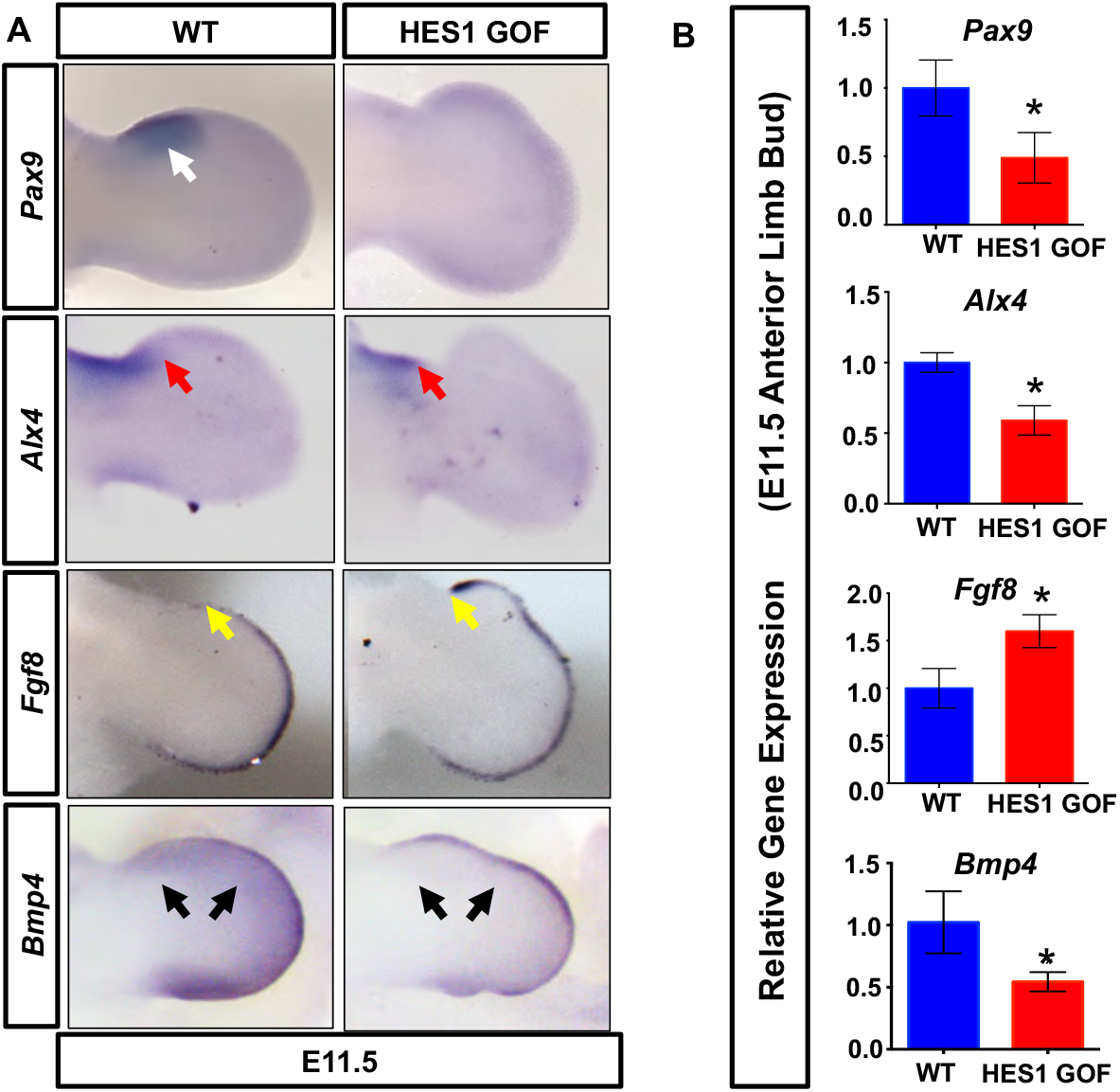
*Hes1* over-expression within limb bud mesenchyme alters the expression of digit patterning factors. **(A)** WISH for *Pax9, Alx4, Fgf8* and *Bmp4* on E11.5 WT and HES1 GOF limb buds. **(B)** qPCR performed on RNA from E11.5 WT and HES1 GOF limb buds (N=4). Asterisks indicate significance with a p-value < 0.05 (Student’s t-test).

To determine whether *Shh* expression and SHH signaling was altered in HES1 GOF mutants, we first performed WISH and qPCR for *Shh* using whole limb buds. HES1 GOF limb buds display both a mild expansion and enhancement in *Shh* expression at E11.5 (**Supp. Fig. 6A,B**). To assess whether the altered *Shh* expression results in enhanced SHH signaling, we assessed SHH target gene (*Ptch1, Gli1,* and *Grem1*) expression and identified no changes when comparing whole limb buds or anterior halves of HES1 GOF and WT limb buds via WISH or qPCR (**Supp. Fig. 6A,B**). Since the processing of GLI3 from its full-length active form (GLI3A) to its repressor form (GLI3R) serves as a readout of SHH activity, we performed Westerns for GLI3 and demonstrate unaltered GLI3R and GLI3A in HES1 GOF limb buds (**Supp. Fig. 6C**). These data indicate that over-expression of *Hes1* within limb bud mesenchyme functions independent or downstream of SHH/GLI3/GREM1 signaling in regulating digit number.

### HES1 functions independent or downstream of SHH/GLI3 and can compensate for *Shh*-deficiency

To determine whether *Hes1* over-expression can compensate for the loss of *Shh* in regulating digit number, we generated HES1 GOF mutant mice in a *Shh* conditional loss-of-function (SHH LOF) background (*Prx1Cre; R26-Hes1*^*f/f*^; *Shh*^*f/f*^). Skeletal analyses demonstrate that *Prx1Cre; R26-Hes1*^*f/f*^; *Shh*^*f/+*^ embryos develop a PPD phenotype similar to HES1 GOF mutants (approx. 6 digits), while *Prx1Cre; Shh*^*f/f*^ mutants develop 1-3 digits as previously indicated (**Fig. 5A,B**)(Zhu and Mackem 2011). Importantly, *Hes1* over-expression within the limb mesenchyme of SHH LOF mice (*Prx1Cre; R26-Hes1*^*f/f*^; *Shh*^*f/f*^) induces the formation of 2-5 digits in nearly 95% of double mutants (**Fig. 5A,B**). Greater than 60% of double mutants develop 3 or more digits, while only 12.5% of SHH LOF mutants develop 3 digits (**Fig. 5A,B**). More than 56% of SHH LOF embryos develop a single digit, while a mere 5-6% of double mutants exhibit one digit (**Fig. 5A,B**). In addition to the partial rescue in digit number, we also observed a rescue of both zeugopod elements (radius and ulna) in ∼10% of double mutants whereas all SHH LOF mutants develop a single zeugopod element resembling the ulna (**Fig. 5A**). Of note, all SHH LOF mice die at or prior to birth, while all *Prx1Cre; R26-Hes1*^*f/f*^; *Shh*^*f/f*^ double mutants generated to date survive to adulthood. These results provide strong evidence that *Hes1* over-expression within *Prx1Cre*-expressing mesenchymal cells is largely capable of compensating for the loss of *Shh* in regulating digit number, patterning of the zeugopod, and survival.

**Figure 5.**
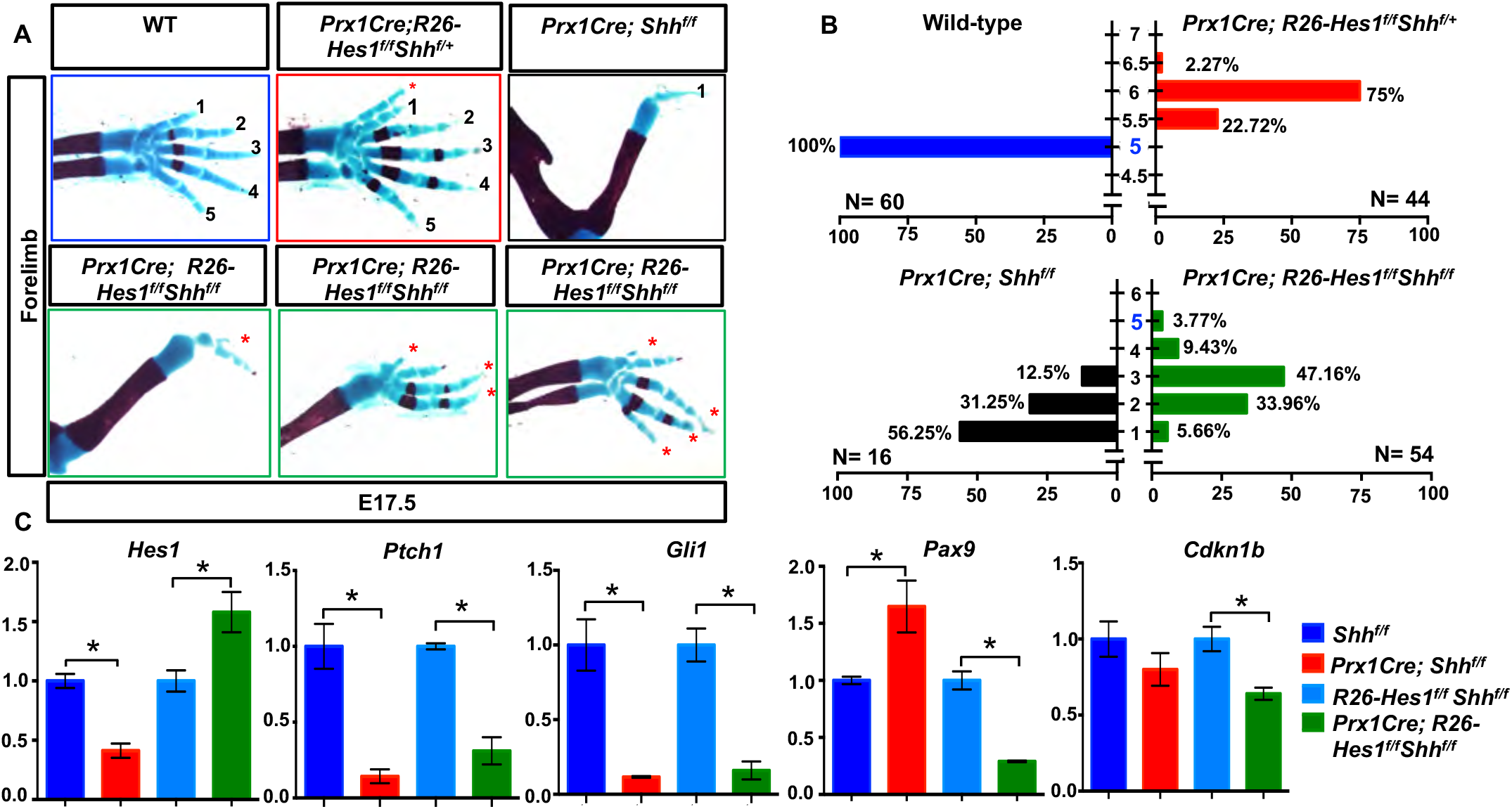
*Hes1* over-expression within limb bud mesenchyme is sufficient to overcome the digit loss observed in SHH LOF limbs. **(A)** Alcian Blue/Alizarin Red staining of WT, *Prx1Cre; R26-Hes1*^*f/f*^; *Shh*^*f/+*^ (HES1 GOF), *Prx1Cre; Shh*^*f/f*^ (SHH LOF), and *Prx1Cre; R26-Hes1*^*f/f*^; *Shh*^*f/f*^ (SHH LOF/HES1 GOF) double mutant forelimbs. Red asterisks indicate HES1-induced digits **(B)** Quantification of digit numbers for WT (n= 60), HES1 GOF (n= 44), SHH LOF (N=16), and SHH LOF/HES1 GOF (n=54) forelimbs (χ^2^ test) (p-value <0.0001). Intervals of 0.5 represent syndactylous digits. **(C)** qPCR for *Hes1, Ptch1, Gli1, Pax9,* and *Cdkn1b* performed on RNA from *Shh*^*f/f*^ (Control), *Prx1Cre; Shh*^*f/f*^ (SHH LOF), *R26-Hes1*^*f/f*^; *Shh*^*f/f*^ (Control), and *Prx1Cre; R26-Hes1*^*f/f*^; *Shh*^*f/f*^ (SHH LOF/HES1 GOF) limb buds at E11.5 (N=4). Asterisks indicate significance with a p-value < 0.05 (Student’s t-test).

To further understand the molecular mechanisms associated with these genetic changes and the corrections to SHH LOF mutant phenotypes by over-expressing *Hes1*, we performed qPCR for *Hes1* and a number of SHH targets, patterning factors, and cell cycle regulators using RNA collected from E11.5 whole limb buds of the genotypes: *Shh*^*f/f*^ (Control), *Prx1Cre; Shh*^*f/f*^ (SHH LOF), *R26-Hes1*^*f/f*^; *Shh*^*f/f*^ (Control), and *Prx1Cre; R26-Hes1*^*f/f*^; *Shh*^*f/f*^ (SHH LOF/HES1 GOF) (**Fig. 5C**). *Hes1* expression is decreased in SHH LOF limb buds, however over-expression of *Hes1* within the limb bud mesenchyme is sufficient to drive *Hes1* expression above both SHH LOF and control levels in SHH LOF/HES1 GOF limb buds (**Fig. 5C**). The SHH target genes, *Ptch1* and *Gli1,* are markedly reduced in SHH LOF limb buds, while over-expression of *Hes1* in the absence or presence of *Shh* is incapable of regulating their expression (**Fig. 5C and Supp. Fig. 6B**). Importantly, *Pax9* exhibits increased expression within SHH LOF limb buds, however over-expression of *Hes1* in the absence of *Shh* dramatically reverses this effect leading to strong suppression of *Pax9* (**Fig. 5C**); also observed in HES1 GOF limb buds (**Fig. 4B**). *Cdkn1b*, a negative regulator of the cell cycle and a direct target of HES1, shows no change in expression when comparing SHH LOF limb buds to controls, however forced expression of *Hes1* in the absence of *Shh* suppresses *Cdkn1b* expression (**Fig. 5C**) similar to that observed in HES1 GOF limb buds (**Fig. 2D**). Therefore, *Hes1* over-expression within the limb mesenchyme can reverse many phenotypic features observed in SHH LOF mutant mice, without directly affecting canonical SHH/GLI3 signaling.

### HES1 and SHH/GLI3 signaling synergize to regulate digit number and gene expression

To study potential cooperative effects of HES1 and SHH/GLI3 signaling in regulating digit number, we developed HES1 GOF mutants in the *Gli3*^*xt/+*^ (GLI3 HET) background (*Prx1Cre; R26-Hes1*^*f/f*^; *Gli3xt*^*/+*^) and appropriate controls. While GLI3 HET and HES1 GOF mice exhibits 5.5 digits (syndactylous 6^th^ digit) or a complete 6 digits, the combination of these alleles induces the formation of more than 6 digits in all *Prx1Cre; R26-Hes1*^*f/f*^; *Gli3*^*xt/+*^ double mutants, thereby producing a more severe PPD phenotype than either single mutant alone (**Fig. 6A,B**).

**Figure 6.**
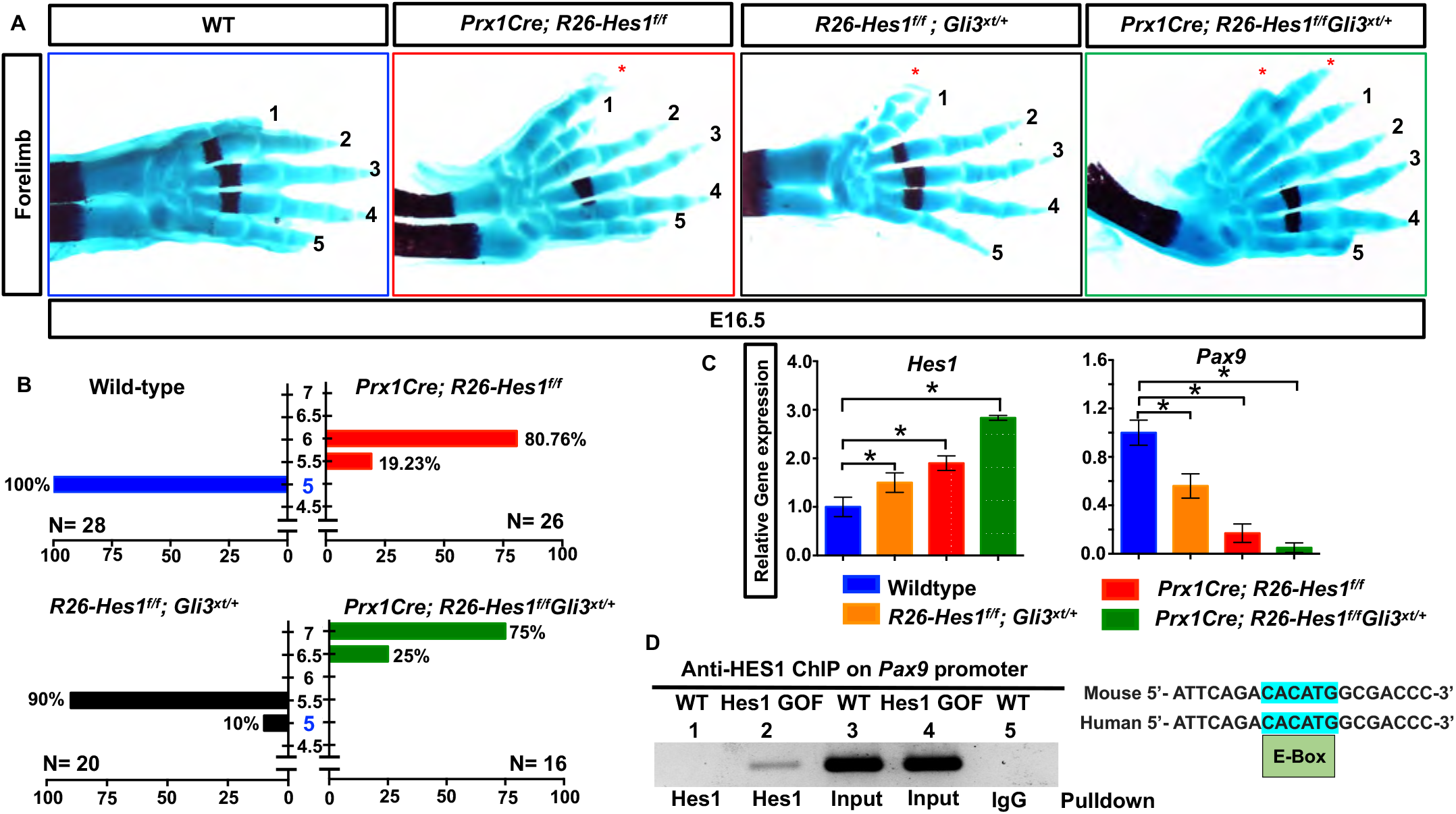
HES1 and SHH/GLI3 signaling cooperatively regulate digit number. **(A)** Alcian Blue/Alizarin Red staining of WT, *Prx1Cre; R26-Hes1*^*f/f*^(HES1 GOF), *R26-Hes1*^*f/f*^; *Gli3*^*xt/+*^ (GLI3 HET), and *Prx1Cre; R26-Hes1*^*f/f*^; *Gli3*^*xt/+*^ (HES1 GOF/GLI3 HET) double mutant forelimbs. Red asterisks indicate extra digits **(B)** Quantification of digit numbers for WT (N=26), HES1 GOF (N=28), GLI3 HET (N= 20), and HES1 GOF/GLI3 HET (N=16) forelimbs (χ^2^ test) (p-value <0.0001). Intervals of 0.5 represent syndactylous digits. **(C)** qPCR for *Hes1* and *Pax9* on RNA from E11.5 WT, *Prx1Cre; R26-Hes1*^*f/f*^(HES1 GOF), *R26-Hes1*^*f/f*^; *Gli3*^*xt/+*^ (GLI3 HET), and *Prx1Cre; R26-Hes1*^*f/f*^; *Gli3*^*xt/+*^ (HES1 GOF/GLI3 HET) double mutant limb buds (N=4). Asterisks indicate significance with a p-value < 0.05 (One-way ANOVA). **(D)** ChIP and PCR amplification of chromatin containing E-Box within *Pax9* promoter (N=3). Lane 1 = amplification of WT chromatin pulled down with anti-HES1; Lane 2 = amplification of Hes1 GOF chromatin pulled down with anti-HES1; Lane 3 = amplification of WT input control chromatin (positive control). Lane 4 = amplification of Hes1 GOF input control chromatin (positive control). Lane 5 = amplification of WT chromatin pulled down with anti-IgG (negative control).

At the molecular level, we also observe an additive effect of these alleles on the expression of specific genes. qPCR utilizing RNA isolated from E11.5 anterior limb buds demonstrates an increase in *Hes1* expression in GLI3 HET limb buds, which was further upregulated in a progressive manner in HES1 GOF and *Prx1Cre; R26-Hes1*^*f/f*^; *Gli3*^*xt/+*^ double mutant limb buds (**Fig. 6C**). Concomitantly, a progressive downregulation of *Pax9* expression occurs while *Hes1* expression increases (**Fig. 6C**). Due to this dynamic inverse relationship and the critical nature of *Pax9* in regulating anterior digit number(Peters et al. 1998), we performed sequence analyses of the *Pax9* promoter and ChIP for HES1. HES1 is capable of binding a specific and conserved E-box within the *Pax9* promoter located approximately 8400bp upstream of the transcriptional start site, however was incapable of binding other non-conserved N-box sequences within the same promoter region of the mouse *Pax9* gene (**Fig. 6D and Supp. Fig. 4C**). These data indicate that SHH/GLI3 and HES1 signals cooperate in regulating digit number and do so in part via the direct HES1-mediated regulation of *Pax9* within the limb bud mesenchyme.

### HES1 functions downstream of SHH/GLI3 signaling and is required for *Gli3*-haploinsufficient PPD

To determine whether HES1 functions downstream of SHH/GLI3 signaling and is required for *Gli3*-haploinsufficient PPD, we generated conditional homozygous deletions of *Hes1* floxed alleles within the limb bud mesenchyme of *Gli3*^*xt/+*^ mutant mice (*Prx1Cre; Hes1^f/f^; Gli3^xt/+^*). Skeletal analyses demonstrate that more than 80% of the *Hes1*^*f/f*^; *Gli3*^*xt/+*^ (GLI3 HET) pups develop a syndactylous polydactyly (5.5 digits) phenotype in this genetic background, while only ∼47% of the *Prx1Cre; Hes1*^*f/f*^; *Gli3*^*xt/+*^ mutant pups develop the same phenotype (**Fig. 7A,B**). These data indicate that HES1 functions downstream of GLI3 and is largely required for the polydactyly phenotype resulting from genetic ablation of a single *Gli3* allele.

**Figure 7.**
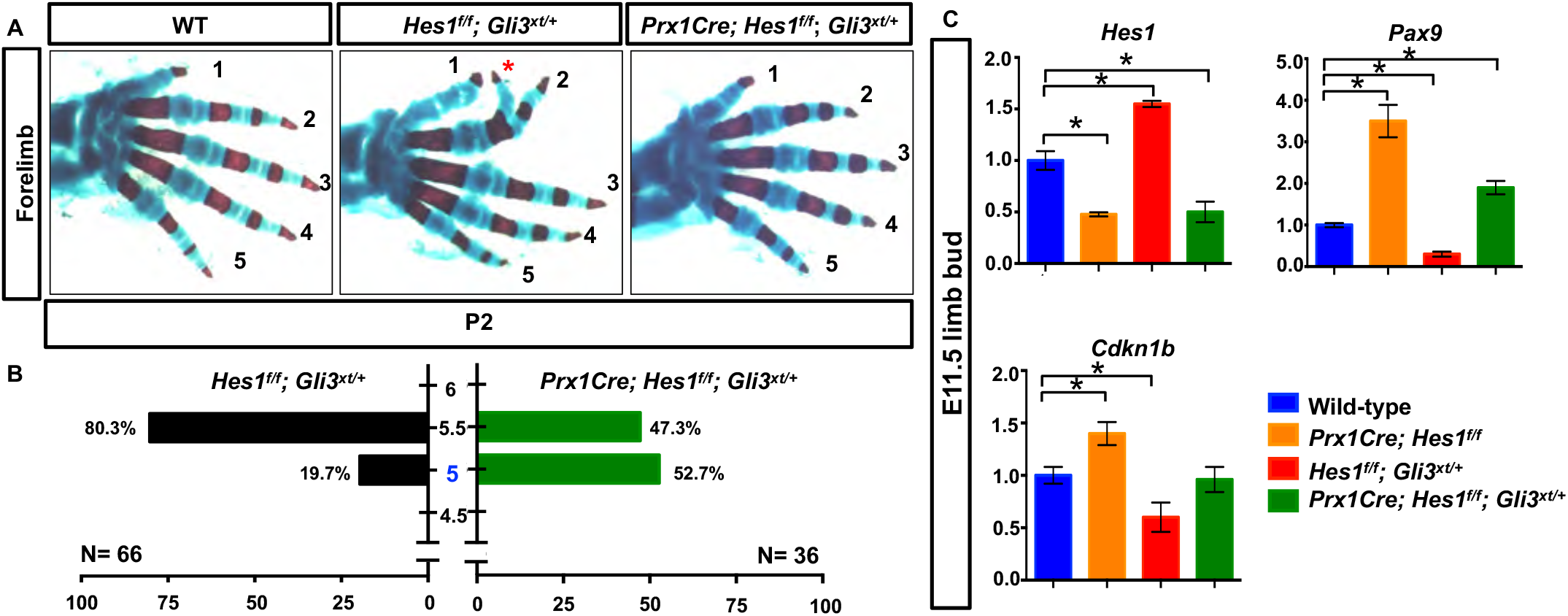
HES1 is a critical effector of SHH-induced PPD. **(A)** Alcian Blue/Alizarin Red staining of WT, *Hes1*^*f/f*^; *Gli3*^*xt/+*^ (GLI3 HET), and *Prx1Cre; Hes1*^*f/f*^; *Gli3*^*xt/+*^ (HES1 LOF/GLI3 HET) double mutant forelimbs. Red asterisk indicates syndactylous digit. **(B)** Quantification of digit numbers for WT, GLI3 HET (N=66) and HES1 LOF/GLI3 HET (N=36) double mutant forelimbs (χ^2^ test) (p-value <0.0001). Intervals of 0.5 represent syndactylous digits. **(C)** qPCR for *Hes1, Pax9,* and *Cdkn1b* on RNA from E11.5 WT, *Prx1Cre; Hes1*^*f/f*^ (HES1 LOF), *Hes1*^*f/f*^; *Gli3*^*xt/+*^ (GLI3 HET), and *Prx1Cre; Hes1*^*f/f*^; *Gli3*^*xt/+*^ (HES1 LOF/GLI3 HET) double mutant limb buds (N=3). Asterisks indicate significance with a p-value < 0.05 (One-way ANOVA).

To further understand the molecular mechanisms associated with the partial genetic rescue of the *Gli3*^*xt/+*^ polydactylous phenotype, we performed qPCR for *Hes1* using RNA collected from E11.5 limb buds of the following genotypes: *Hes1*^*f/f*^ (WT controls), *Prx1Cre; Hes1*^*f/f*^ (HES1 LOF), *R26-Hes1*^*f/f*^; *Gli3*^*xt/+*^ (GLI3 HET), and *Prx1Cre; Hes1*^*f/f*^; *Gli3*^*xt/+*^ (HES1 LOF/GLI3 HET) (**Fig. 7C**). *Hes1* expression decreases in both HES1 LOF and HES1 LOF/GLI3 HET limb buds, while it increases in GLI3 HET limb buds (**Fig. 7C; also Fig. 6C**). These data suggest that *Hes1* is a target of SHH/GLI3 signaling in the regulation of digit number. Previous studies identified GLI factors as potential direct regulators of *Hes1(Vokes et al. 2008; Wall et al. 2009)*. Therefore, we performed sequence analyses of the *Hes1* promoter and ChIP utilizing anti-GLI2 and anti-GLI3 antibodies to precipitate chromatin isolated from E11.5 WT limb buds. These data demonstrate that GLI3 and GLI2 are each capable of binding two different consensus sequences within the *Hes1* promoter located approximately 146bp and 8150bp upstream of the transcriptional start site (**Supp. Fig. 7**). Collectively, these data indicate a negative regulation of *Hes1* by GLI3R within the limb bud mesenchyme during limb development.

Since we identified *Pax9* and *Cdkn1b* as potentially important direct HES1 gene targets during HES1-induced PPD, we assessed the expression of these genes in WT, HES1 LOF, GLI3 HET, and HES1 LOF/GLI3 HET limb buds. The expression of both *Pax9* and *Cdkn1b* are upregulated in HES1 LOF limb buds and downregulated in GLI3 HET limb buds, however HES1 LOF/GLI3 HET rescued mutant limb buds exhibit *Pax9* and *Cdkn1b* expression levels similar to or slightly higher than WT controls (**Fig. 7C**). These data suggest that SHH/GLI regulation of digit number is dependent on HES1 and the HES1 transcriptional targets, *Cdkn1b* and *Pax9*, which control both proliferation of the limb mesenchyme and anterior boundaries of digit formation.

## DISCUSSION

Cooperative regulation of mesenchymal cell proliferation and chondrogenic differentiation is critical in regulating the size of the limb field and digit number. As chondrogenesis within the limb skeleton proceeds in a proximal to distal direction, the apical zone mesenchyme of the autopod is influenced by signals from the ZPA (SHH) and AER (FGFs)(Riddle et al. 1993; Laufer et al. 1994; Chiang et al. 2001). FGFs are required for mesenchymal cell survival and to reinforce *Shh* expression within the ZPA(Sun et al. 2002). SHH in turn regulates mesenchymal cell proliferation by relieving GLI3 repression, resulting in the induction of *Cdk6* and *Ccnd1,* among other positive regulators of the cell cycle(Vokes et al. 2008; Lopez-Rios et al. 2012; Lewandowski et al. 2015). Additionally, SHH/GLI signaling induces *Grem1,* which functions to inhibit BMP-induced cell cycle exit and chondrogenesis to maintain apical zone mesenchymal cells in a primitive state until SHH/GLI signaling is down-regulated and BMP signals initiate the condensation of SOX9-expressing mesenchymal progenitors(Panman et al. 2006; Lopez-Rios et al. 2012). *Gli3* deficiency results in enhanced *Cdk6, Ccnd1,* and *Grem1* expression, which promotes expansion of the limb field via increased mesenchymal cell proliferation, delayed BMP-induced cell cycle exit, and delayed chondrogenesis. This allows for the development of additional mesenchymal condensations within the expanded autopod resulting in polydactyly(Lopez-Rios et al. 2012). Highlighting the importance of the tight control of mesenchymal proliferation and differentiation in regulating digit number, *Gli3*-deficient polydactyly is partially overcome via the conditional removal of *Cdk6* or a single copy of *Grem1(Lopez-Rios et al. 2012)* or exacerbated by deletion of one or more *Bmp4* alleles(Dunn et al. 1997; Lopez-Rios et al. 2012). Further demonstrating the exquisite sensitivity of this program, conditional deletion of *Bmp4*(Selever et al. 2004) alone or overexpression of *Grem1*(Norrie et al. 2014) within limb bud mesenchyme delays mesenchymal cell cycle exit and the onset of chondrogenesis resulting in a polydactyly phenotype resembling that of *Gli3*-deficient mice. Here we describe a novel and requisite role for HES1 in regulating digit number in the developing autopod downstream of SHH/GLI3 signaling. Our data indicate that GLI3R directly suppresses *Hes1* expression in the anterior mesenchyme, while active SHH/GLI signaling promotes *Hes1* expression in the distal posterior mesenchyme of the developing limb bud. Mechanistically, HES1 inhibits chondrogenesis and stimulates mesenchymal cell proliferation by directly repressing chondrogenic genes (*e.g. Col2a1*) and negative regulators of mesenchymal cell proliferation (*e.g. Cdkn1b/*p27), which appears to be distinct from the positive and direct regulation of cell cycle inducers (*e.g. Cdk2, Cdk4, Cdk6*) and the activation of a potent inhibitor of BMP-induced cell cycle exit and chondrogenesis (*e.g. Grem1*) via SHH/GLI3. Therefore, HES1 mediates unique aspects of SHH/GLI signaling to expand the limb field to the appropriate size via the regulation of both mesenchymal cell proliferation and the onset of chondrogenesis.

In addition to BMP signaling and SHH-induced regulators of the cell cycle, anterior expressed genes such as *Alx4* and *Pax9* are under the control of SHH/GLI signaling and aid in establishing anterior boundaries for autopod expansion(Peters et al. 1998; Kuijper et al. 2005; McGlinn et al. 2005; Panman et al. 2005). ALX4 restricts the expression of *Shh* and *5’Hoxd* genes to the distal posterior mesenchyme(Kuijper et al. 2005; Panman et al. 2005), and inhibits cell proliferation in some contexts(Shi et al. 2017). PAX9 is a critical regulator of A-P patterning and boundary formation in multiple tissues including the limb bud, odontogenic mesenchyme, and the palatal mesenchyme(Peters et al. 1998; Zhou et al. 2013; Sivakamasundari et al. 2017). In both of the latter instances, PAX9 regulates the expression of *Bmp4* and *Msx1*, while PAX9 and MSX1 cooperatively and directly regulate the transcriptional activation of *Bmp4(Kong et al. 2011; Zhou et al. 2013; Sivakamasundari et al. 2017)*. Both *Alx4* and *Pax9* are reduced or absent in the limb mesenchyme of *Gli3*^*xt/+*^ or *Gli3*^*xt/xt*^ polydactylous mice and enhanced throughout the mesenchyme in *Shh*^*−/−*^ limb buds(Kuijper et al. 2005; McGlinn et al. 2005; Panman et al. 2005). Interestingly, *Bmp4* expression correlates with GLI3R levels and is subsequently reduced in the anterior mesenchyme of *Gli3*^*xt/xt*^ limb buds and expressed throughout the mesenchyme of *Shh*^*−/−*^ limb buds(Bastida et al. 2004). Since genetic ablation of *Pax9* and conditional inactivation of *Msx1* and *Msx2* within the limb mesenchyme results in preaxial polydactyly phenotypes resembling *Gli3*^*xt/+*^ mice(Peters et al. 1998; Bensoussan-Trigano et al. 2011), it is likely that *Bmp4* expression within the anterior limb mesenchyme is transcriptionally regulated via PAX9-MSX1 in a manner similar to that observed in the odontogenic and palatal mesenchyme(Kong et al. 2011; Zhou et al. 2013). While PAX9 regulation of *Bmp4* within the anterior mesenchyme provides a likely mechanism by which PAX9 establishes an anterior boundary for digit development via regulation of cell cycle exit and chondrogenesis, the precise molecular mechanism by which SHH/GLI signaling regulates the expression of *Pax9* has remained elusive. Our data indicates a critical role for HES1 downstream of SHH/GLI3 signaling to directly regulate *Pax9* expression. In *Gli3*-deficient mice, as in HES1 GOF mice, HES1 expression is observed broadly within the posterior and anterior limb bud mesenchyme resulting in the direct suppression of *Pax9* that in turn reduces the expression of *Bmp4*. Conversely, in the *Shh*-deficient limb bud, GLI3R is expressed throughout the limb mesenchyme suppressing *Hes1* expression which leads to the subsequent posterior expansion of both *Pax9* and *Bmp4;* contributing factors to *Shh*-deficient digit reductions. Collectively, all of our data are consistent with, and fit into the greater framework of, digit number regulation as demonstrated at the anatomic and molecular levels by numerous genetic models, including *Gli3*-deficiency(Litingtung et al. 2002; Lopez-Rios et al. 2012)*, Pax9*-deficiency(Peters et al. 1998)*, Msx1/Msx2*-deficiency(Bensoussan-Trigano et al. 2011), *Bmp4*-deficiency(Dunn et al. 1997; Selever et al. 2004), *Grem1*-over-expression(Norrie et al. 2014), and *Shh*-deficiency(Litingtung et al. 2002; Zhu et al. 2008; Zhu and Mackem 2011), among others. Our findings further provide novel insights into SHH/GLI regulation of *Hes1*, as well as, HES1 direct regulation of *Cdkn1b* and *Pax9* and their roles in regulating digit number, *Gli3*-deficient polydactyly, and *Shh*-deficient digit reductions. The SHH/GLI/HES1 signaling axis identified here functions in parallel with the SHH/GLI/GREMLIN axis previously identified as being critical for normal digit development(Lopez-Rios et al. 2012), highlighting the complexity of integrated signals necessary to coordinate proper pentadactylous digit formation within the developing autopod (**Supp. Fig. 8**).

While we have identified HES1 as a new and critical regulator of digit number downstream of SHH/GLI signaling, some questions remain regarding this complex signaling regulation of digit development. *Hes1* is an established target gene of the NOTCH signaling pathway and the NOTCH ligand, *Jagged1*, and NOTCH target gene, *Hey1,* are also regulated via SHH/GLI signaling, raising questions as to whether the NOTCH pathway itself functions downstream or in parallel to SHH/GLI-mediated control of digit number(McGlinn et al. 2005; Vokes et al. 2008; Wall et al. 2009; Lewandowski et al. 2015). Similar to our HES1 GOF mutant mice and *Gli3*-deficient mice, overexpression of the NOTCH intracellular domain (NICD) within the limb bud mesenchyme (*Prx1Cre; R26-NICD*^*f/+*^) results in enhanced mesenchymal cell proliferation, however completely inhibits chondrogenesis as compared to a temporary delay in chondrogenesis observed in HES1 GOF and *Gli3*-deficient limb buds(Hui and Joyner 1993; Dong et al. 2010). Alternatively, conditional deletion of *Rbpjk* floxed alleles within the limb mesenchyme (*Prx1Cre; Rbpjk*^*f/f*^) accelerates chondrogenesis and only reduces skeletal element or digit size as compared to *Shh*-deficient digit number reductions(Kraus et al. 2001; Dong et al. 2010). Interestingly, autosomal dominant forms of Adams-Oliver Syndrome (OMIM 614814, 616028, 616589) caused by germline loss-of-function mutations in *RBPjk*, *NOTCH1*, and the NOTCH ligand, *DLL4*, commonly present with abnormalities of the hands and feet including fused digits (syndactyly), severe shortening of digits (brachydactyly), and/or complete loss of digits (oligodactyly). These most severe types of digit reductions are likely not observed in our conditional NOTCH pathway loss-of-function mice (*Prx1Cre; Rbpjk*^*f/f*^ or *Prx1Cre; Hes1*^*f/f*^ as examples)(Dong et al. 2010; Rutkowski et al. 2016) for a number of reasons, including the timing by which the *Prx1Cre* transgene induces gene deletion or the limited expression of the transgene to only the skeletogenic mesenchyme. Similar differences between germline deletions and conditional deletions induced by the *Prx1Cre* transgene can be highlighted by the differential digit reductions observed in *Prx1Cre; Shh*^*f/f*^ and *Shh*^*−/−*^ mutant mice(Kraus et al. 2001; Zhu and Mackem 2011). SHH/GLI signaling may indeed regulate *Hes1* in both NOTCH-dependent and -independent manners, therefore future studies will be required to tease apart whether NOTCH signaling itself is required for *Gli3*-deficient polydactyly or whether *Shh*-deficient digit restrictions can be overcome by some form of NOTCH activation.

## MATERIALS AND METHODS

### Mouse strains

The *Prx1Cre* mouse line was obtained from JAX laboratory and previously described(Logan et al. 2002). The *R26-Hes1*^*f/f*^ and *Hes1*^*f/f*^ mouse lines were a generous gift from Dr. Ryoichiro Kageyama (Kyoto University)(Kobayashi et al. 2009). The *Gli3*^*xt/xt*^ (Maynard et al. 2002) and *Shh*^*f/f*^ (Lewis et al. 2001) mutant mice were also obtained from JAX labs. Embryos and mice were harvested at E10.5, E11.5, E12.5, E16.5, E18.5 and 2 months. All animal work was approved by both the Duke University and University of Rochester Institutional Animal Care and Use Committees (IACUC).

### Whole mount skeletal staining and in situ hybridization

Whole embryo skeletal staining for digit number analysis was performed using the protocol as previously described(Rigueur and Lyons 2014). To look at spatial-temporal expression of various genes we performed WISH on E11.5 limb buds using the previously described protocol(Rutkowski et al. 2014).

### RNA isolation and qPCR

Whole and anterior limb buds were harvested in cold 1× PBS and flash frozen using liquid nitrogen. After genotypes were obtained, mutant and wild type limb buds were pooled and homogenized in Trizol (Invitrogen) using a 25g needle. The RNA was then precipitated using 1-bromo-3-choloropropane (MRC). The aqueous layer was then separated and washed with 70% ethanol and purified using the RNeasy Plus Mini Kit (Qiagen). The RNA was transcribed into cDNA and qPCR was performed using methods described (Dong et al. 2010). Mouse specific primers sequences are listed in Supplemental Table 1.

### Chromatin immunoprecipitation (ChIP) assay

The ChIP assay was performed on wild type and Hes1 GOF E11.5 limb buds using the MAGnify Chromatin Immunoprecipitation system (Invitrogen) using the protocol described (Rutkowski et al. 2016). The HES1 antibody was a generous donation from Dr. Ryoichiro Kageyama (Kyoto University). The antibody was used at a concentration of 5 μg per reaction. Data analysis was performed using PCR primers specifically designed flanking the conserved E-box or N-box sequences within the *Cdkn1b*, *Col2a1,* and *Pax9* promoter or enhancer regions. Primers are listed in Supplemental Table 1.

### Western blots

Whole and anterior limb buds at E11.5 were isolated from wild type and Hes1 GOF embryos and lysed in RIPA buffer (Invitrogen) containing protease inhibitors. The lysate was then centrifuged, the supernatant containing the protein was obtained and quantified using a BCA system. Protein (15µg) was separated using NuPAGE Novex 8% and 10% Bis-Tris pre-cast gels (Invitrogen) and was transferred to a PVDF membrane using the BIORAD system. Antibodies against HES1 (1:1000; Cell Signaling), GLI3 (1:1000; R&D systems); P27 (1:1000; BD Biosciences), CCND1 (1:1000; Cell Signaling), LMNA (1:1000; Abcam), ACTIN (1:2000; Sigma-Aldrich), and TUBULIN (1:1000; Cell Signaling) were used with the appropriate secondary antibodies. Antibody information is located in Supplemental Table 2.

### Immunohistochemical and histological staining

Embryos were harvested in cold 1× PBS, fixed in 4% paraformaldehyde (PFA), the limbs were dissected and then hand processed. Limbs were embedded in OCT for frozen sectioning and paraffin for standard microtomy and IHC or histologic staining. Frozen sections were cut at 10μm, while paraffin sections were cut at 5μm. To analyze the general cellular morphology, standard histological staining using ABH–OG was performed. IHC was performed on paraffin sections using the VectaStain ABC kits and developed with ImmPACT DAB (Vector Labs). Primary antibodies against the following proteins were used for IHC analyses: ACAN (1:200; Chemicon), COL2A1 (1:100; Thermo Scientific), SOX9 (1:100; Santa Cruz Biotechnology), and PH3 (1:200; Cell Signaling). Immunohistochemistry for BrdU (Invitrogen) was performed as previously described (Dong et al. 2010).

### Statistical analysis

Statistical analyses were performed using two-tailed Student’s *t*-test and one way ANOVA; a *P*-value <0.05 was considered significant (denoted by * in the figures).

## AUTHOR CONTRIBUTIONS

Conception and study design: MJH and DS; Study conduct: DS, AJM, JL, and AL; Analysis and interpretation of the data: DS, AJM, JL, AL, CMK, and MJH; Drafting and/or editing manuscript: DS, CMK, and MJH; Approving final version of the manuscript: DS, AJM, JL, AL, CMK, and MJH.

## ACKNOWLEDGEMENTS

This work was supported in part by the following United States National Institute of Health grants: R01 grants (AR057022, AR063071, and AR071722 to MJH), and departmental funds from the Department of Orthopaedic Surgery at Duke University School of Medicine to MJH and CMK. We thank Dr. Ryoichiro Kageyama (Kyoto University) for providing important *Hes1* floxed and *R26-Hes1* mouse strains, as well as, HES1 antibodies.

## SUPPLEMENTAL FIGURES

**Supplemental Fig 1.**
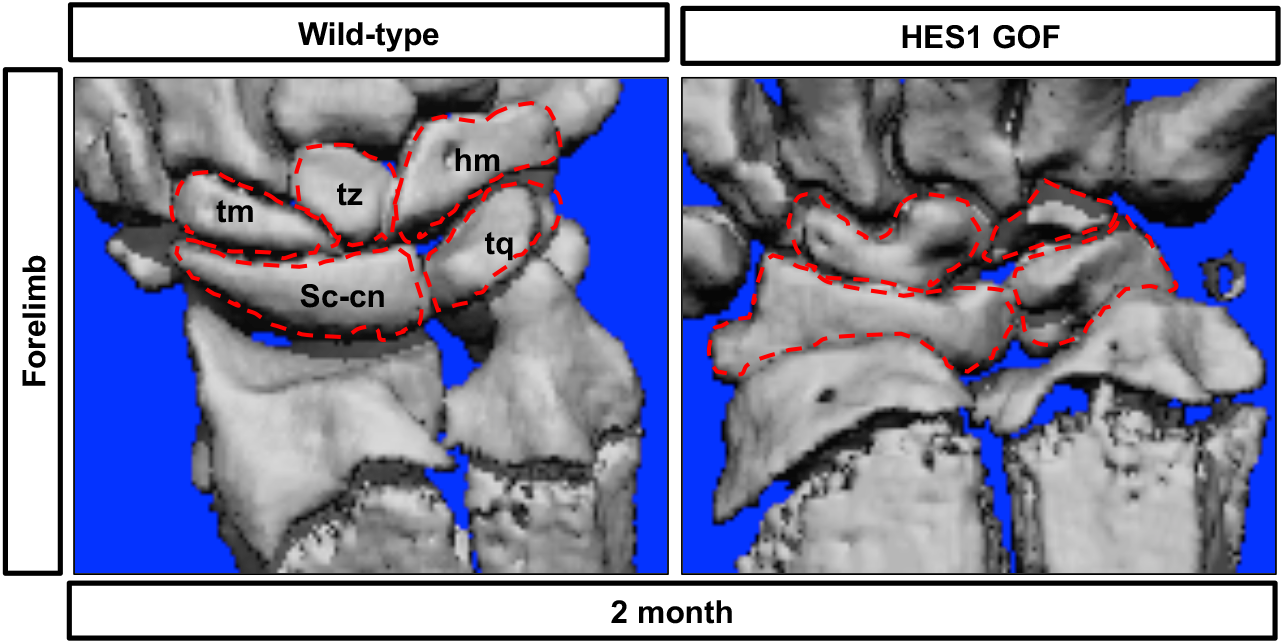
*Hes1* over-expression within limb bud mesenchyme results in defects in carpal joint formation. MicroCT images of the carpal bones from WT and HES1 GOF mice at 2-month of age. hm, hamate; sc-cn, scaphoid-centrale; tq, triquetral; tm, trapezium; tz, trapezoid. Red dashed lines indicate location of carpal joints.

**Supplemental Fig 2.**
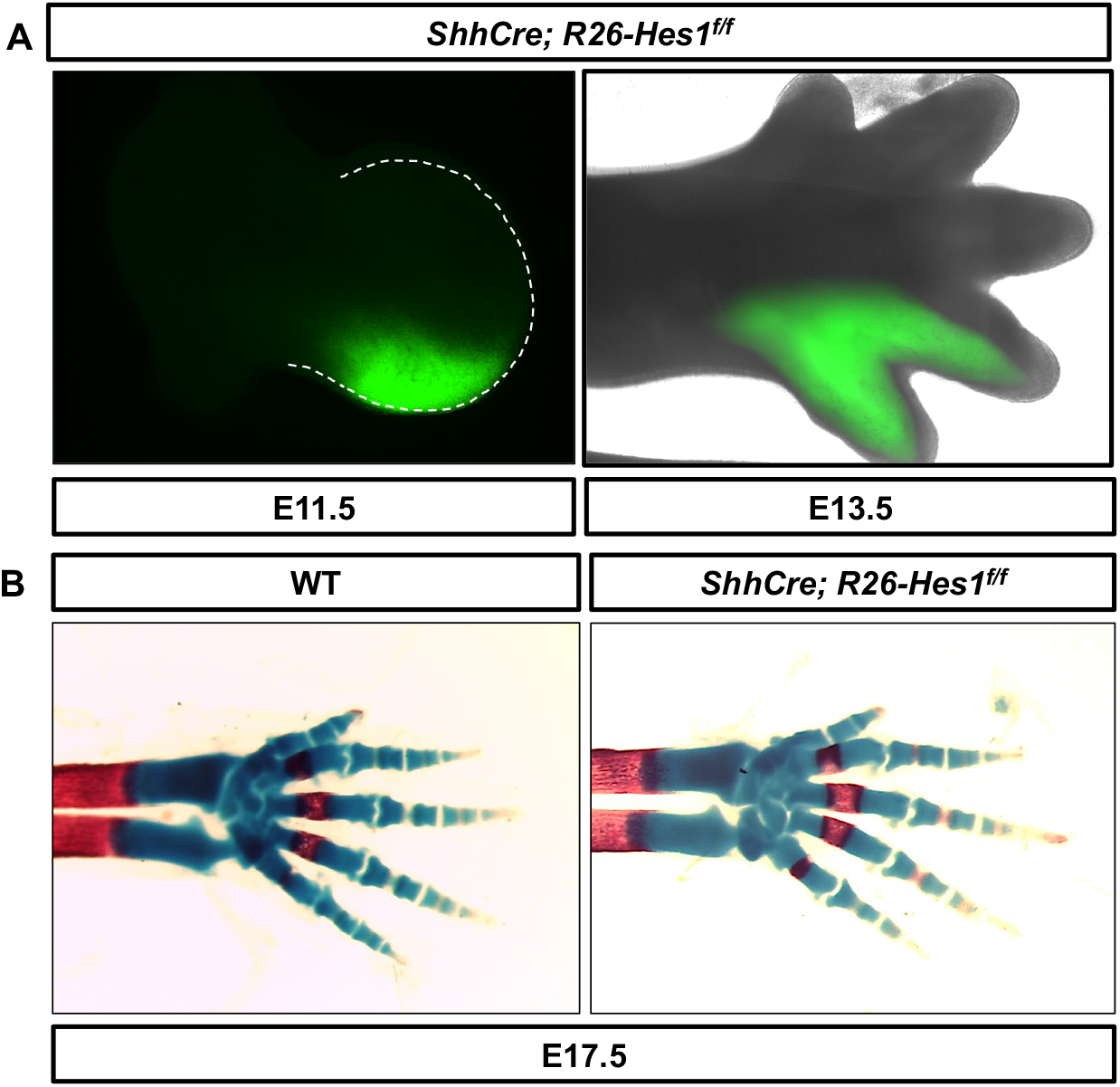
*Hes1* over-expression within the posterior limb bud mesenchyme does not induce PPD. **(A)** GFP fluorescence from a *ShhCre; R26-Hes1*^*f/f*^ limb bud at E11.5 and forelimb at E13.5 (N= 6). R26-Hes1 floxed allele contains and IRES-GFP labeling *ShhCre* expressing cells and their descendants. **(B)** Alcian Blue/Alizarin Red staining of WT and *ShhCre; R26-Hes1*^*f/f*^ mutant forelimbs at E17.5 (N=12).

**Supplemental Fig 3.**
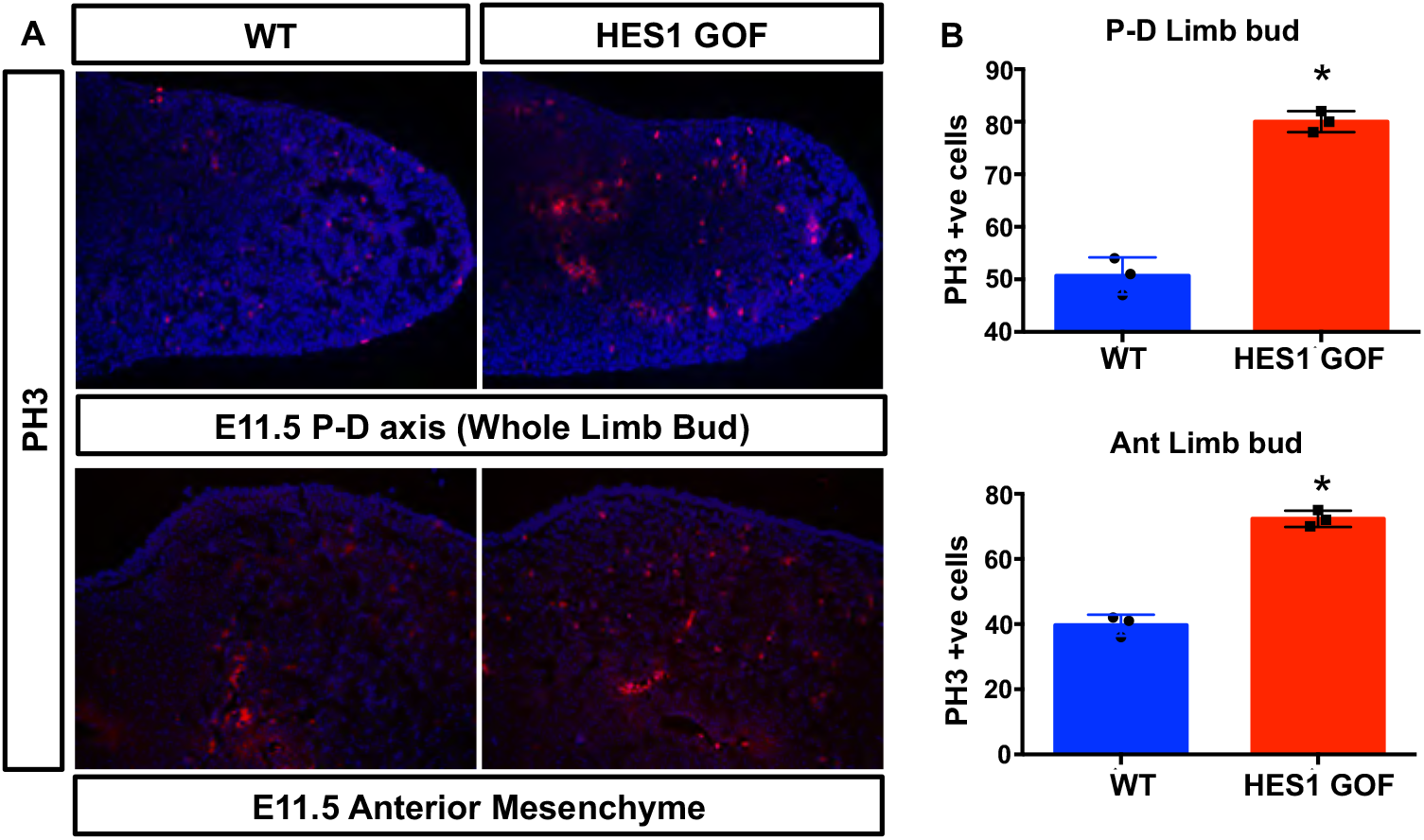
*Hes1* over-expression within limb bud mesenchyme induces phospho-histone H3. **(A)** Immunoflourescence for PH3 along the proximal-distal (P-D) axis and anterior mesenchyme of E11.5 HES1 GOF and WT limb buds **(B)** Quantification of PH3 positive cells along the P-D axis and anterior mesenchyme (N=3). Asterisks indicate significance with a p-value < 0.05 (Student’s t-test).

**Supplemental Fig 4.**
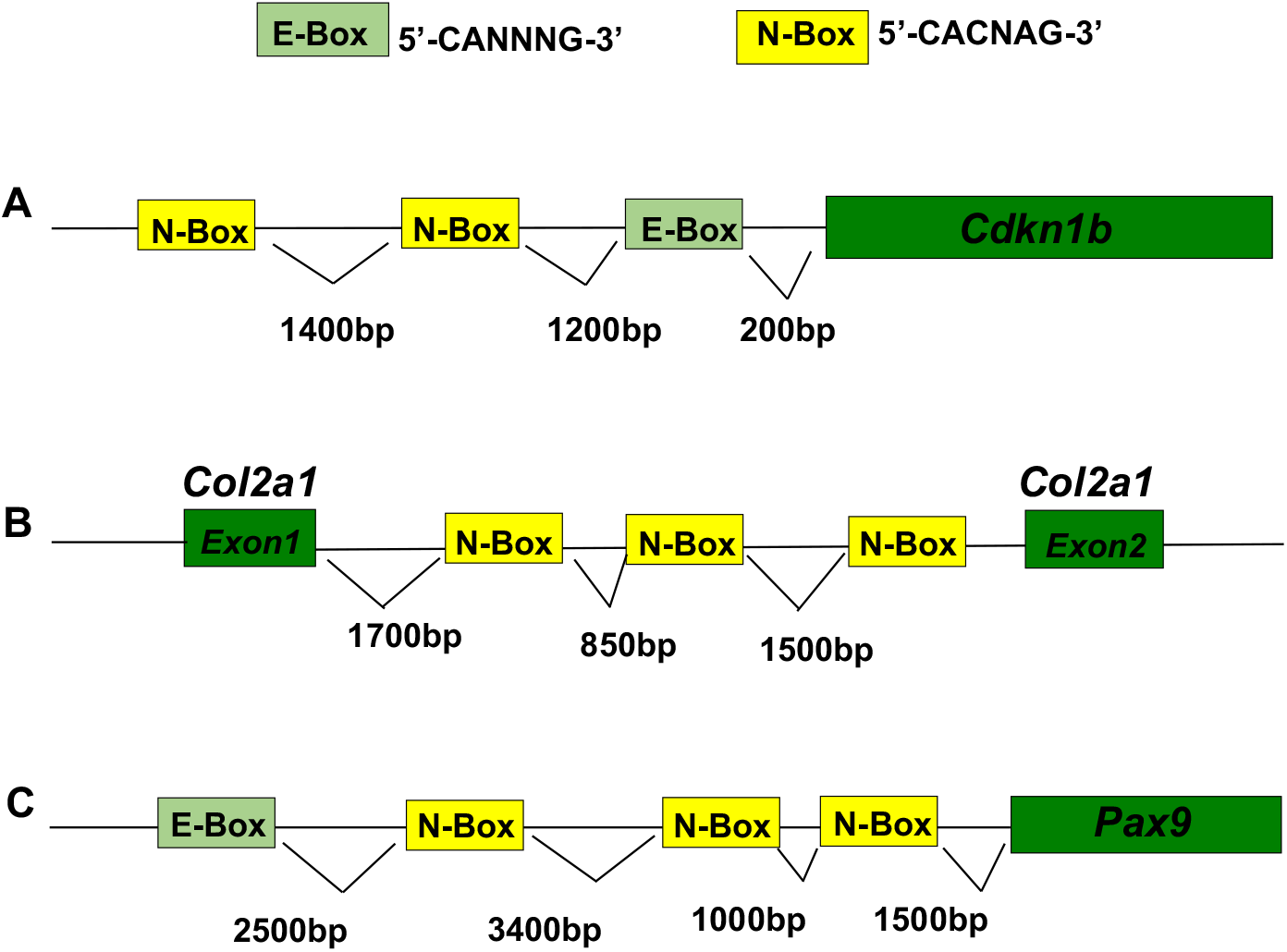
HES1 binding sites within the *Cdkn1b, Col2a1*, and *Pax9* promoter/enhancer regions. **(A)** Schematic of potential HES1 binding sites within the *Cdkn1b* promoter. **(B)** Schematic of potential HES1 binding sites within the *Col2a1* enhancer region located between exons 1 and 2. **(C)** Schematic of potential HES1 binding sites within the *Pax9* promoter. Approximate distances in base pairs (bp) upstream of transcriptional start sites or downstream from exons are indicated for each potential HES1 binding site.

**Supplemental Fig 5.**
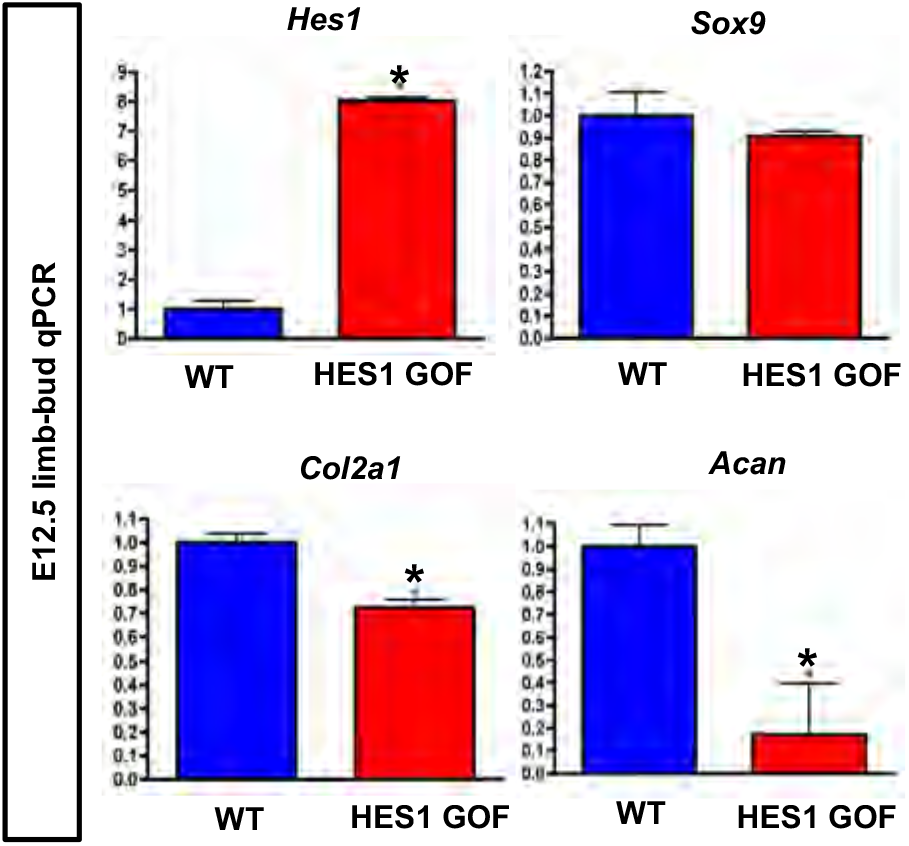
*Hes1* over-expression within limb bud mesenchyme suppresses chondrogenic gene expression at E12.5. qPCR for *Hes1, Sox9, Col2a1,* and *Acan* on RNA isolated from WT and HES1 GOF E12.5 whole limb buds (N=4). Asterisks indicate significance with a p-value < 0.05 (Student’s t-test).

**Supplemental Fig 6.**
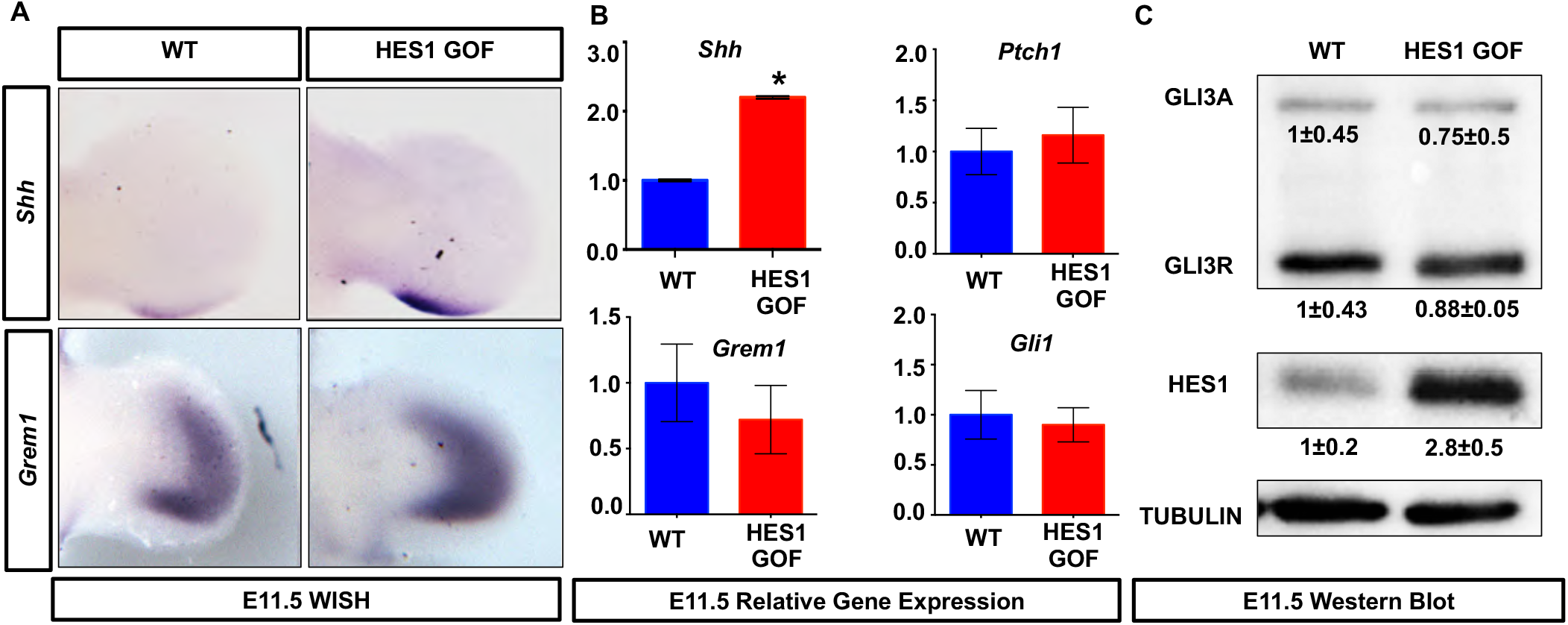
*Hes1* over-expression within limb bud mesenchyme enhances *Shh* expression, but not SHH signaling. **(A)** WISH for *Shh* and *Grem1* on E11.5 WT and HES1 GOF limb buds. **(B)** qPCR for *Shh, Ptch1, Grem1,* and *Gli1* on RNA isolated from E11.5 WT and HES1 GOF limb buds (N=4). Asterisks indicate significance with a p-value < 0.05 (Student’s t-test). **(C)** Western blots performed on protein extracted from E11.5 WT and HES1 GOF limb buds for GLI3, HES1, and TUBULIN (N=4). Quantification represented as Fold change± SD relative to WT.

**Supplemental Fig 7.**
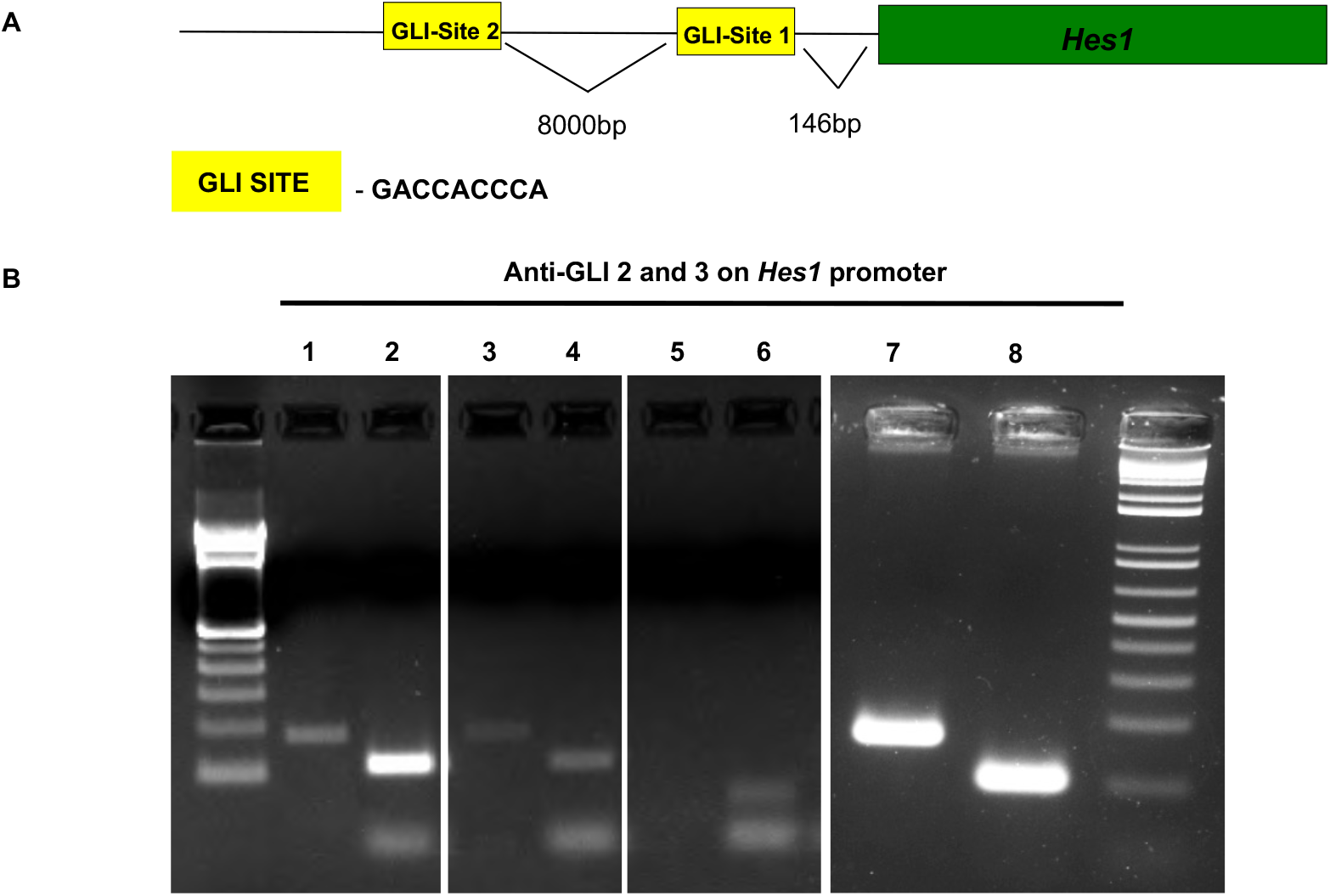
GLI3 binds specific regions of the *Hes1* promoter. **(A)** Schematic of potential GLI binding sites within the *Hes1* promoter. **(B)** ChIP and PCR amplification of chromatin containing GLI binding sites within *Hes1* promoter (N=3). Lane 1 = amplification of Site 1 using WT chromatin pulled down with anti-GLI3; Lane 2 = amplification of Site 2 using WT chromatin pulled down with anti-GLI3; Lane 3 = amplification of Site 1 using WT chromatin pulled down with anti-GLI2; Lane 4 = amplification of Site 2 using WT chromatin pulled down with anti-GLI2; Lane 5 = amplification of WT chromatin pulled down with anti-IgG (negative control); Lane 6: amplification of site 2 using WT chromatin pulled down with anti-IgG (negative control). Lane 7 = amplification of Site 1 using input control chromatin (positive control); Lane 8 = amplification of Site 2 using input control chromatin (positive control).

**Supplemental Fig 8.**
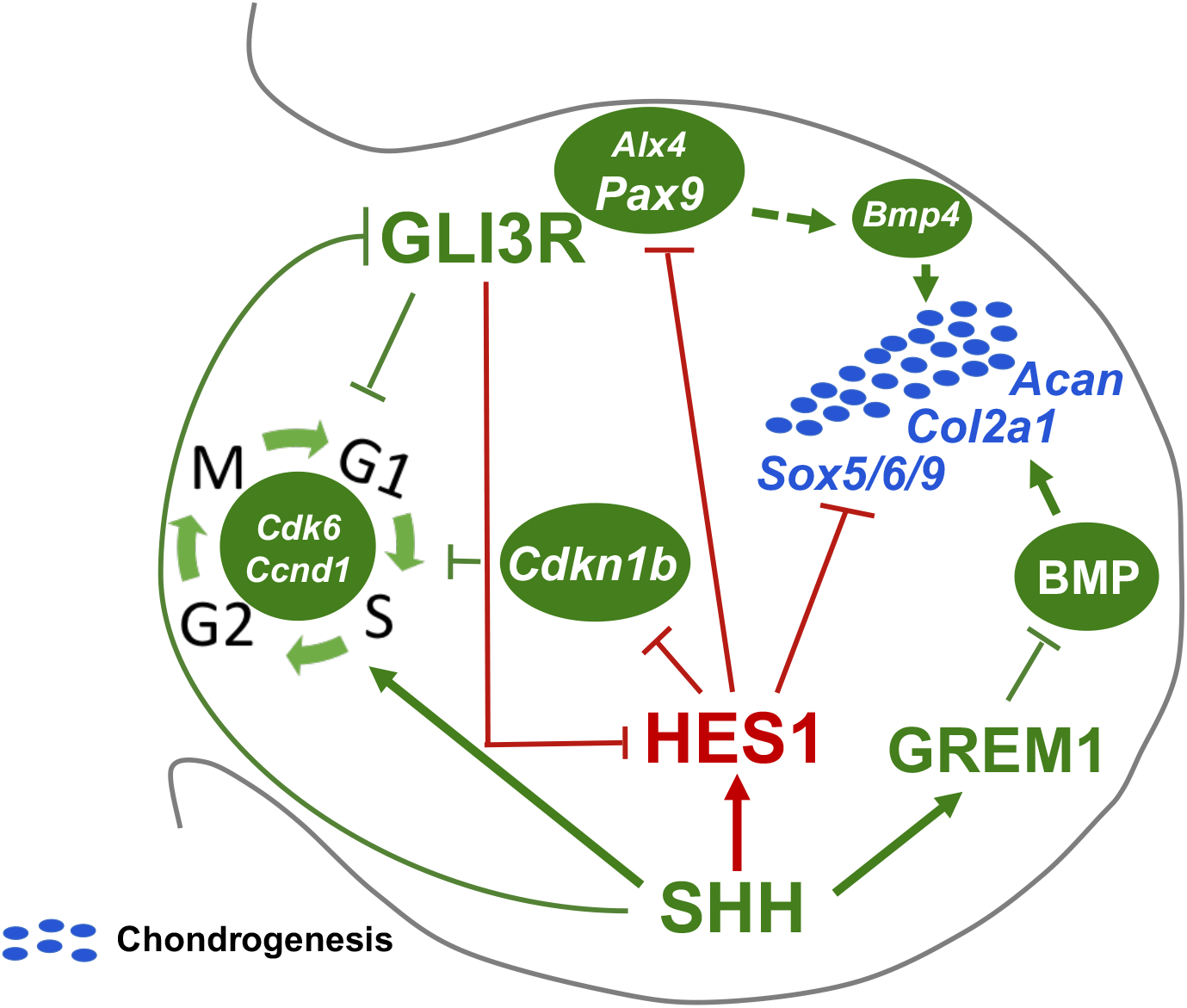
Proposed model for SHH/GLI3/HES1 regulation of digit number. HES1 functions downstream of SHH/GLI3 signaling to regulate digit number via: 1) a HES1-mediated transcriptional regulation of *Cdkn1b* expression to coordinate mesenchymal cell proliferation, 2) a HES1-mediated transcriptional regulation of *Pax9* to establish anterior boundaries of digit chondrogenesis, and 3) a HES1-mediated and BMP/GREM1 signaling independent transcriptional regulation that coordinates the timing of digit chondrogenesis. Red lines and arrows indicate novel findings in the SHH/GLI3/HES1-mediated regulation of digit number.

## SUPPLEMENTAL TABLES

**Supplemental Table 1.**
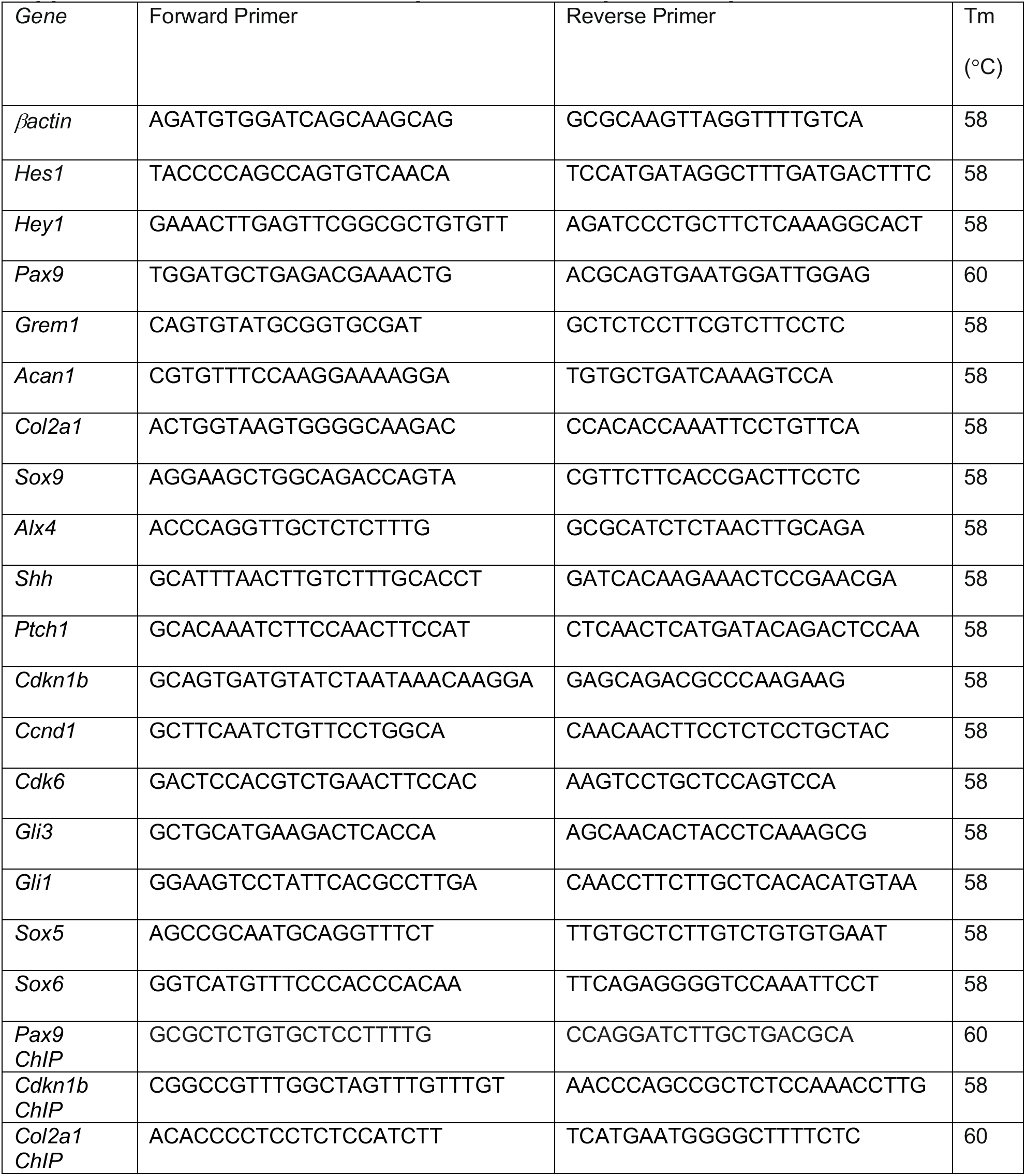
Real-time qPCR and ChIP primer sequences.

**Supplemental Table 2.** List of antibodies.

| **Antibody** | **Company and Catalog** | **Concentration** |
| --- | --- | --- |
| Hes1 | Cell Signaling 11988 | 1:1000 |
| Gli3 | R&D systems AF3690 | 1:1000 |
| Gli2 | Abcam ab26056 | 1:1000 |
| P27 | BD biosciences 610241 | 1:1000 |
| Cyclin D1 | Cell Signaling 2926 | 1:1000 |
| Actin | Sigma Aldrich 4970 | 1:2000 |
| LaminB1 | Abcam Ab16048 | 1:2000 |
| PH3 | Cell Signaling 9701 | 1:200 |
| Col2a1 | Thermo Scientific MS235-p | 1:200 |
| BrdU | Invitrogen 93-3943 | 1:1000 |
| HRP Anti-Mouse | Cell Signaling 7076S | 1:2000 |
| HRP Anti- Rabbit | Cell Signaling 7074S | 1:2000 |
| HRP Anti- Goat | R&D systems HA7017 | 1:1000 |
| Goat Anti-Rabbit 568 | Invitrogen A-11011 | 1:200 |
| Biotinylated Horse anti-mouse | Vector Labs BA-2000 | 1:1000 |
| Biotinylated Goat anti-rabbit | Vector Labs BA-1000 | 1:1000 |

